# The hippocampus as an epistemic forager: When curiosity and reward jointly steer exploration and hippocampal replay

**DOI:** 10.1101/2025.10.31.685837

**Authors:** Erik Németh, Augustin Chartouny, Krisztina Jedlovszky, Ismael T. Freire, Mehdi Khamassi

## Abstract

Awake hippocampal replay is a widely studied phenomenon wherein special neurons of the hippocampus encoding spatial locations – place cells – show a sequential reactivation during periods of immobility, often representing trajectories to or from reward locations, as observed in foraging rodents. Several computational models have been proposed to explain how this phenomenon could contribute to memory consolidation and action planning. However, certain aspects of the mechanism behind hippocampal replay remain unclear, such as why reactivation is biased towards not only reward sites but also decision points. Here, we propose that both expected reward (satisfying hunger) and expected information gain (satisfying curiosity) contribute to determine the priority of events to be replayed. To test this, we present the Epistemic Replay Algorithm (ERA), which bridges reinforcement learning and active inference into a single computational model. We evaluate ERA in five experiments spanning three maze types: linear maze, non-stationary maze and double T-maze. Our results first showcase that more curious agents explore more thoroughly and adapt faster to changing environments while still being capable of exploiting optimal rewards. Further, we find that the ERA model accounts for a larger number of hippocampal replay properties compared to non-curious models, including (i) a broad-to-specific progression of hippocampal replay events; (ii) symmetric replay around decision points; and (iii) the preferential pausing at both reward sites and decision points. To conclude, we derive new predictions to further test the model and discuss its implications compared to alternative accounts.

## 1. Introduction

Why does the hippocampus sometimes replay trajectories that do not lead directly to reward? Hippocampal replay is a neural phenomenon observed in mammals such as rodents, bats, non-human primates and humans [1, 2, 3, 4], that plays a key role in spatial cognition and memory consolidation. During active exploration of an environment, hippocampal place cells [5] fire in sequences that correspond to the animal’s trajectory through space [6, 7]. Afterwards, during wakeful rest or slow-wave sleep, these same sequences are reactivated – or “replayed” – at compressed timescales, often coordinated with sharp-wave ripple oscillations in the hippocampus [8, 9]. Replay can occur in the same order as the original experience (forward replay) or in reverse order (reverse replay), which is thought to contribute to learning from recent outcomes [10]. This process is believed to strengthen neural representations of space, linking experiences across time, and supporting spatial memory [11, 12], planning and decision-making by allowing the animal to simulate possible future trajectories [13, 14]. Yet, awake replay often occurs at decision points [15] and in familiar areas, even when no immediate reward is at stake [16]. What drives this “curious” behavior?

Existing computational frameworks provide important but incomplete answers. State-of-the-art models treat awake hippocampal replay as selective, utility-driven sampling that supports planning and credit assignment [17, 18]. In a similar fashion, in reinforcement learning, “Dyna”-style and prioritized schemes formalize which past transitions should be replayed by their expected improvement to future reward [19, 17, 20]. These normative accounts explain forward, reverse and remote replay patterns and their sensitivity to reward changes, aligning replay content with the value of computation during deliberation.

Predictive-map theories based on the successor representation complement this view by casting the hippocampus as learning state-to-state transition frequencies [21]. Whether the hippocampus is involved in learning local transition probabilities as in such models, or whether it approximates a global transition structure of a full “cognitive map”, replay in all these models operates over the established transition probabilities to enable rapid revaluation when rewards or policies change. In the active-inference framework, replay is framed as generative sampling of candidate trajectories from a hierarchical world model to minimize expected free energy [22, 23, 24]. This formulation naturally accounts for goal-dependent sequences that reduce uncertainty and support compositional inference during learning and navigation. Across these accounts, replay converges onto a unified computational role as a model-based evaluation mechanism that selectively prioritizes trajectories with maximal expected utility (whether defined by reward revaluation or information gain) to drive adaptive decision-making. Nevertheless, to our knowledge, while the effect of expected reward gain on both task performance and hippocampal replay patterns has been studied [17, 19], expected information gain has only been considered in terms of its effect on task performance [24]. In other words, it is not yet clear what specific neural activity patterns would be predicted by a replay mechanism prioritizing *both* reward and information gain.

The groundwork of seminal replay algorithms was laid by Peng and Williams [25] and Moore and Atkeson [26], publishing their independent research in the very same year. What these *prioritized sweeping* -type RL algorithms have in common is (i) a selection of high-impact experience stored in a “replay buffer”; (ii) the simulation of counterfactual high-impact experiences via the transition model, and their addition to the same buffer; and (iii) RL-style inference over the highest-priority states until the buffer is empty or convergence is achieved within some tolerance range. Despite the simplicity and ensuing popularity of the prioritized sweeping framework, several variants of this algorithm exist (Tab. 1). Nevertheless, while these RL-based architectures can readily explain many of the key aspects of hippocampal reactivation, the question of why non-rewarding trajectories are often preferentially replayed [27, 28] remains debated.

**Table 1:**
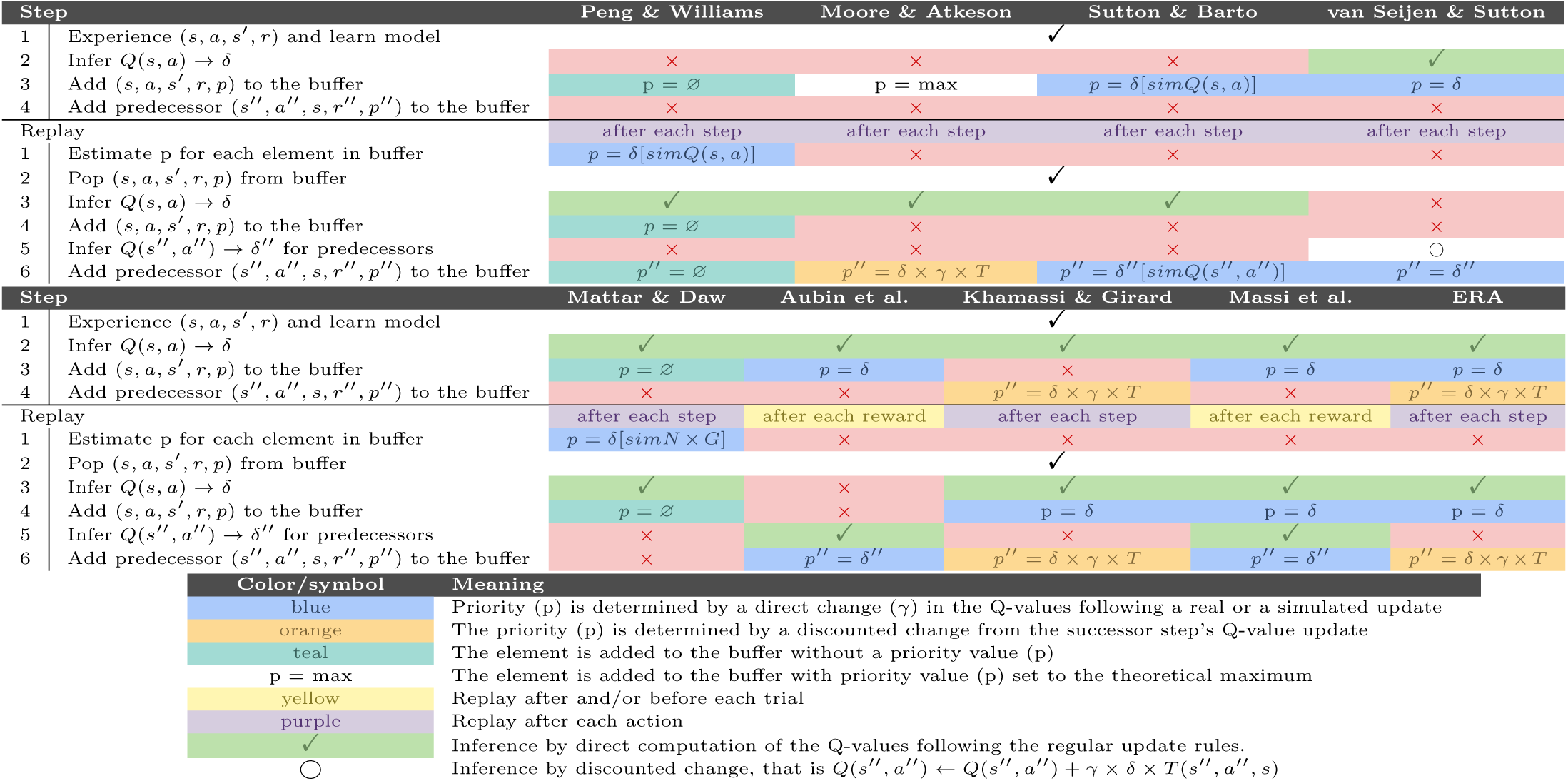
A comparative analysis of different solutions of prioritized sweeping. The algorithm is decomposed into 4 in-real-life and 6 virtual (simulated) steps. The methods compared are Peng and Williams [25], Moore and Atkeson [26], Sutton and Barto [29], Van Seijen and Sutton [30], Mattar and Daw [17] (which Antonov and Dayan [31] are basing themselves on, see Sec. 4 for further detail), Aubin et al. [32], Khamassi and Girard [20], Massi et al. [18] and the ERA algorithm proposed in this paper. *δ* stands for a difference after Q-value update, the nature of which is indicated in brackets if necessary. *sim* means that an update step is computed (simulated) but the update is not applied to the Q-values. *N × G* stands for *Need* and *Gain*, two terms further explained in Sec. 4.

It has been shown that curiosity can contribute to memory formation and retention in humans [33, 34], and models of curiosity have been equally proposed as a natural solution to the exploration-exploitation dilemma central to active inference [35]. As such, it stands to reason that it directly or indirectly impacts hippocampal replay. Accordingly, we hypothesize that a drive towards expected information gain (or curiosity) may play an important role in structuring said process. Not only should immediate information constitute a prompt for reactivation and learning, but agents may seek to plan a sequence of actions through known areas in order to reach unknown loci where they expect high information gain. If this is the case, then the informational content of hippocampal replay at decision points should reflect not only future expected rewards but also future expected information gain. In this way, uninformed, stochastic exploration could be complemented or even replaced by *directed*, informed exploration based on an estimate of discounted information gain [24]. This leads to the novel hypothesis that the epistemic function of the mammalian hippocampus is to use the learned cognitive map of the world to infer the potentially reducible, long-term discounted expected uncertainty with respect to the internal model (as opposed to instantaneous, or one-step ahead information gain). The former could be learned through surprise signals experienced in the world, leading to information gains projected across the entirety of the cognitive map.

In order to test this hypothesis, we propose a new computational model, which we call Epistemic Replay Algorithm (ERA), which bridges model-based reinforcement learning and active inference framework by handling multiple-dimensional utility functions. In particular, we test here the model with 2 dimensions: (1) food reward that animals can acquire in a maze and (2) information gain about food reward uncertainty. The latter constitutes *epistemic rewards*, similarly to the epistemic term used in active inference models [22]. Importantly, ERA enables the estimation of long-term delayed uncertainty: Since our model treats information gains as reward dimensions (in line with the “common currency” hypothesis [36, 37]) on which the Bell-man equation can be applied, it thus enables to learn an expected value *Q^d^* of any reward dimension *d* that approximates the discounted sum of future rewards according to this dimension: 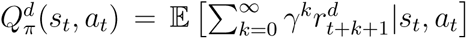 where *π* is the optimal policy with respect to need *d*, *s_t_* and *a_t_* are the current state and action respectively, 0 *< γ <* 1 is temporal discount factor, and *r^d^* is the reward corresponding to dimension *d*. So, when *d* represents the epistemic dimension, such as an information gain in the world model, the theoretical framework of RL enables propagating this epistemic reward to all states of the task through learning and model-based inference, so that *Q^d^* represents the expected future information gain in the distal states of the task. This extends existing computational models, including active inference-based solutions, which to our knowledge were focused on local uncertainty (stimulus uncertainty, state uncertainty, reward uncertainty, etc.).

We perform a series of numerical simulations comprising five experiments across several mazes with diverse topologies and dynamic properties. We show that agents driven by both reward and information gain (i) explore more efficiently, (ii) adapt more rapidly to environmental changes, and (iii) reproduce key features of awake hippocampal replay observed in rodents, including initiation and reward bias, preferential deliberation in decision points (or, more precisely, branching points, see Sec. 3.6), and the symmetric replay of outcomes irrespective of rewards (for further details, see Tab. 2). We show these results stem from a smooth, dynamic, and efficient balance of exploration and exploitation; wherein epistemic exploration – *i.e.*, exploration directed towards delayed expected information gains – is triggered when encountering surprise signals and task changes, followed by a naturally increasing exploitation due to the reduction of uncertainty, until reaching optimal performance asymptotes. From a neurobiological point of view, the simulations presented hereafter show that we can account for a number of properties of awake hippocampal replay, including its increase after task changes and introductions of novel events, its prevailing occurrence at reward sites and decision-(or branching-) points, and its increase with the number of state visits (Tab. 2). Our model simulations also yield novel predictions for neuroscience, suggesting that replay should be particularly prominent in regions of expected information gain, and consequently its structure might resemble more of that of a breadth-first graph search.

**Table 2:** Features of awake hippocampal replay reported in the neuroscience literature and successfully reproduced by the ERA algorithm. The findings marked in bold cannot be (or can only partially be) reproduced by classical, non-curious prioritized sweeping algorithms such as [18].

| General topic | Finding | References |
| --- | --- | --- |
| Replay locations | Initiation bias | [9, 10, 38] |
|  | Reward bias | [39, 40, 41, 6, 42, 43] |
|  | **Preferential stopping in decision points** | [44, 13, 15] |
| Symmetries | Reward apparition and extinction both result in replay | [45, 20] |
|  | **Replay sweeps are not selective towards rewarded side** | [13] |
| Temporal dynamics | Replay is proportional to time spent in given place field | [46] |
|  | More replay in novel environment and after a reward change | [10, 47] |
|  | Replay decreases with exposure | [48] |
|  | **Replay goes from broad (non-specific) to local (specific)** | [17] |

## 2. Materials & Methods

### 2.1 General presentation of the ERA

The Epistemic Replay Algorithm (ERA) is a model-based reinforcement learning algorithm using a multi-dimensional Q-table alongside an infinite replay buffer (Fig. 1).

**Figure 1:**
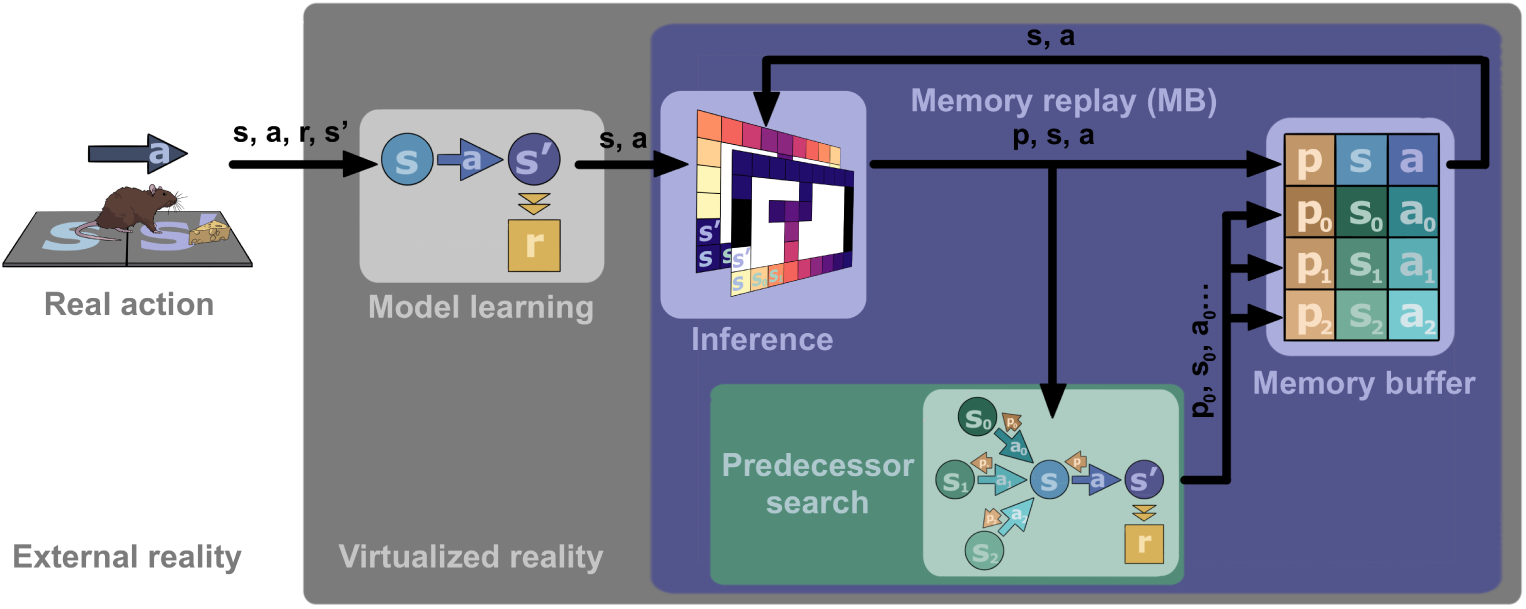
The Epistemic Replay Algorithm (ERA). After each step the agent updates its own world model, which is then used for updating the value of the corresponding state-action values in the two-dimensional Q-table. This transition is stored in the prioritized memory buffer, alongside its predecessor states according to the world model. Then, given a sufficiently high surprise, the top elements of the buffer are replayed.

Observation in the environment are represented by starting states *s ∈ S*, actions *a ∈ A*, arrival states *s′ ∈ S* and food rewards *r^f^ ∈* ℝ. Each transition is stored in an *n*-step transition- and reward history function, which are used to calculate a running average of the transition probabilities *P* (*s′|s, a*) and the expected reward E(*r^f^*). These are stored in the transition function *T* (*s, a, s′*) *∈* [0, 1] and the reward function *R^f^* (*s, a*) *∈* ℝ respectively, which constitute the world model.

To model curiosity as a drive towards minimizing model uncertainty, the reward function is extended by a second dimension, namely the expected information gain (or epistemic reward) *R^i^*(*s, a*) *∈ ℝ*. This quantity is computed as the absolute change in the uncertainty of the world model after its update. Consequently, we denote *R*(*s, a*) = [*R^f^* (*s, a*)*, R^i^*(*s, a*)] *∈* ℝ^2^ as the full, two-dimensional reward-function.

Q-values (expected discounted rewards [29]) are then computed via the the Bellmann-equation:

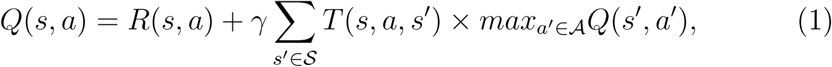

where *γ ∈* [0, 1[ is the discount factor controlling the horizon of the agent. Just as the reward-function, the Q-table also consists of two dimensions *Q*(*s, a*) = [*Q^f^* (*s, a*)*, Q^i^*(*s, a*)] *∈ ℝ*^2^, one food-related and one information-related.

While the Bellmann-update is classically iterated over the entire state space and until convergence, targeted replay – and most notably prioritized sweeping [25, 26] – allows for updating the Q-values where the largest change is expected. This computation is gradually propagated towards more distal regions in the state space, until said change decreases below a threshold *θ ∈* R. Consequently, model updates are efficient, quick, and are only triggered upon encountering surprise.

### 2.2 The experimental setup

In order to reproduce patterns observed in the hippocampal activity in rodents, we opted for modeling a spatial navigation task. Three different types of mazes were proposed across five tasks, each one modeled as a two-dimensional grid world, where states *s* corresponded to physical location, with actions *a ∈ {north, east, south, west, stay}*.

Food rewards *r^f^* were delivered upon the selection of a pre-defined action in a given state, following a Bernoulli distribution with *q ∈* [0, 1] reward delivery probability. Usually, the rewarded action was chosen to be “stay”, with *q* = 1. The agent was allowed to exploit the rewarded action in a repeated fashion until the end of the episode, where each episode was defined via the maximum number of actions taken by the agent. At the end of each episode, the agent was automatically moved back to the start location.

Deviations from this setup will be mentioned in the corresponding sections. Most notably, in almost all cases we chose to slightly restrict the action space in a realistic fashion (e.g. forbidding to choose to bump into the wall of the maze). Furthermore, to test for stochastic rewards, *q* = 0.5 was used in Sec. 3.2; and in order to reproduce previous findings in the double-T maze – where, in the original experiments, rats were prevented from turning back after engaging in a corridor – in Sec. 3.5 we restricted the agent’s motion drastically, making almost all the maze unidirectional (except for two decision points). In this setup, the “stay” action was removed and reward was delivered upon reaching the rewarded state. Given the nature of the maze, the agent in this framework would inevitably return to the start location, thus episodes were defined based on its arrival instead of a fix duration.

### 2.3 Implementation of the ERA model

Epistemic rewards were computed as the absolute change in the uncertainty in the world model after its update, i.e. either in the the (food) reward function *R^f^* (*s, a*) (which transiently re-increases when the reward location is changed) or in the transition-function (which transiently re-increases when a new obstacle is introduced in the environment). In the former case the change of uncertainty was formalized as the difference in the variance of a beta-function fit onto the n-step reward history, a formalism adapted to Bernoulli-distributed food rewards [49] (Sec. 3.1-3.6); and in the latter we opted for the difference in the entropy of the distribution *T_t_*(*s, a,* s′) = *P_t_*(*s′|s, a*) and *T_t−_*_1_(*s, a, s′*) over all *s′ ∈ S* (Sec. 3.7). In either case, the difference was computed upon each model update step, so that both dimensions of *R*(*s, a*) were refreshed at the same time. The length of the time window over which reward history was integrated was selected to be *n* = 2 for most experiments; and *n* = 10 for experiments where it was deemed important to visualize the slow adaptation to stochastic change.

#### Algorithm 1 Predecessor search

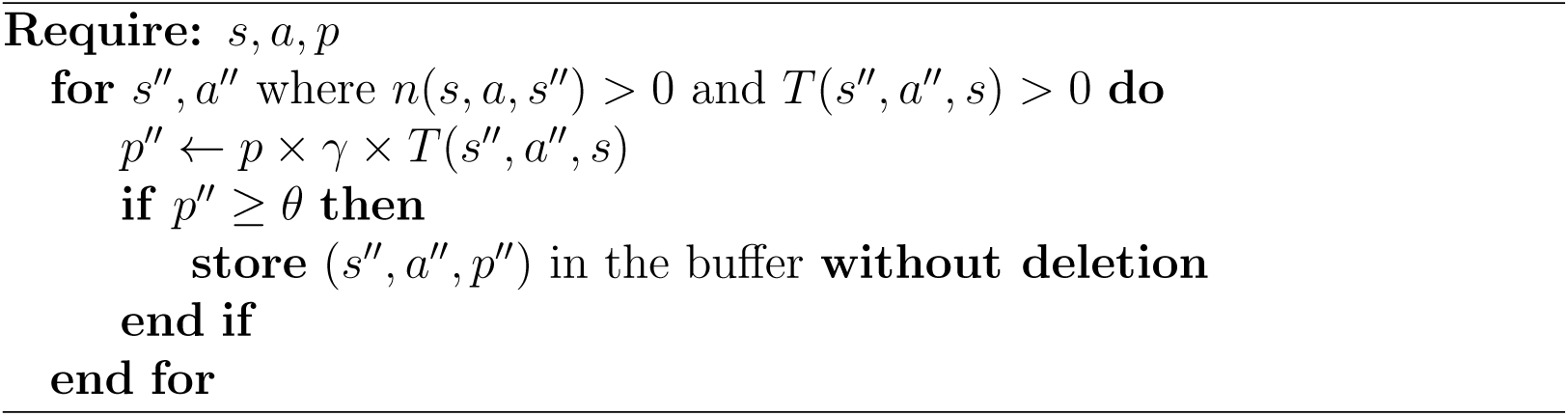

#### Algorithm 2 Memory replay

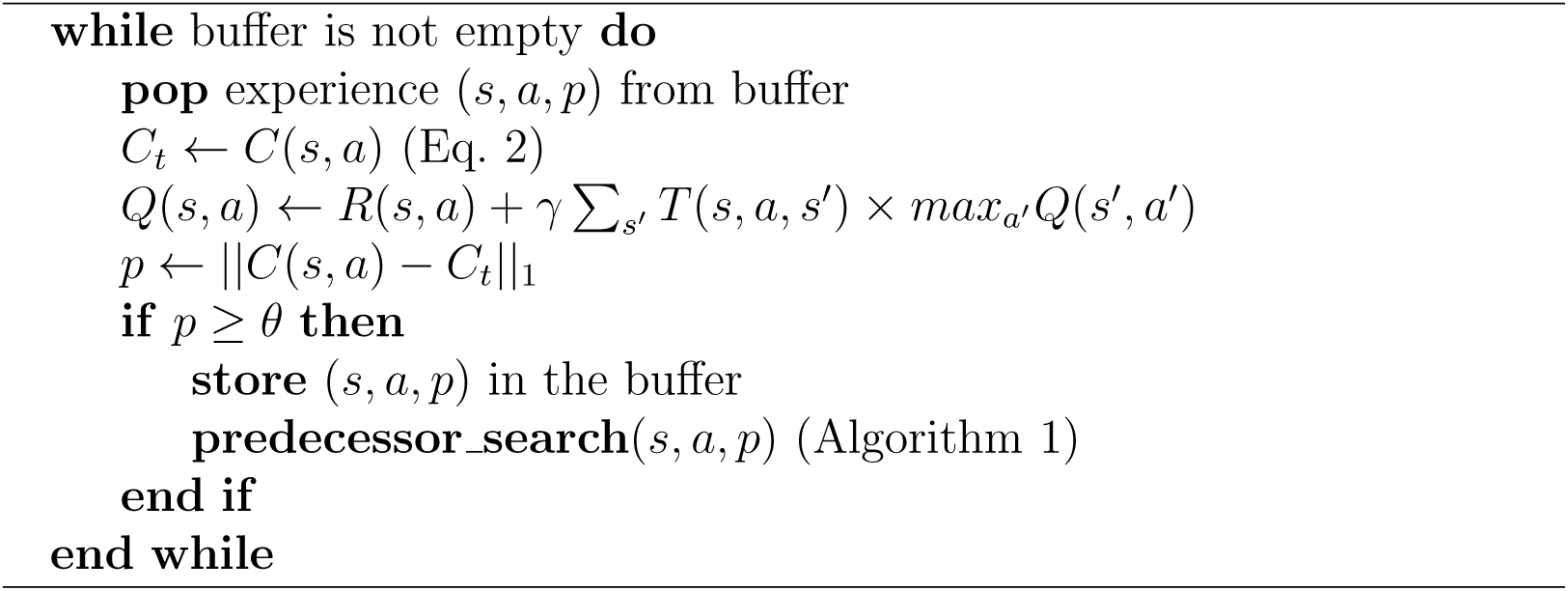

Both in case of the *R*(*s, a*) and the *Q*(*s, a*) functions, the first (food-related) dimension was initialized to zero; while the second (information-related) was initialized to its theoretical maximum, mirroring the maximal uncertainty present in the environment.

#### Algorithm 3 Epistemic Replay Algorithm

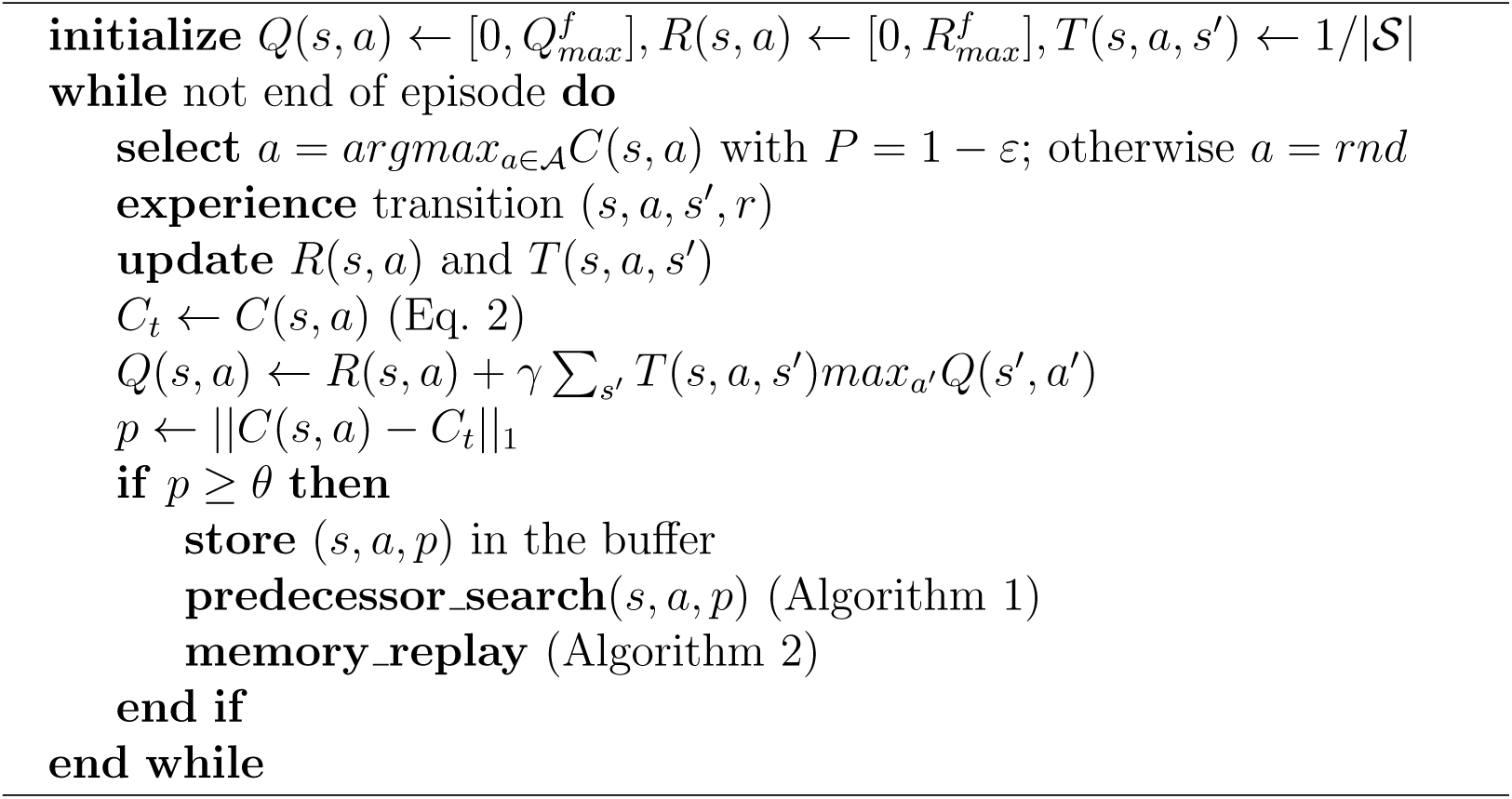

Replay was realized using an infinite memory buffer, storing observations in a prioritized queue, with priority *p*(*s, a*) = ||Δ*C*(*s, a*)||_1_ = ||*C_t_*(*s, a*) *− C_t−_*_1_(*s, a*)||_1_, where

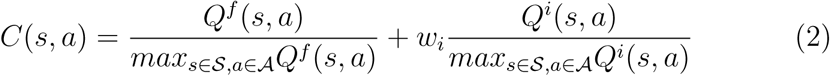

Here *w_i_*is the weight of the information-related component, and is used to control the impact of curiosity on the agent.

Upon an observation with priority *p ≥ θ*, the corresponding (*s, a, p*) triplet was added to the prioritized buffer, the agent suspended its motion, and a predecessor search was initiated (Alg. 1). During this latter, replay priority was back-propagated across the world model, and if the discounted priority exceeded a threshold, i.e. *p′′*(*s′′, a′′*) = *p*(*s, a*) *× γ × T* (*s′′, a′′, s*) *> θ*, the triplet (*s′′, a′′, p′′*) was added to the buffer; or if already present, its priority was updated. These updates were only allowed to promote elements of the buffer, as opposed to demoting or erasing them.

Following the predecessor search, the top element of the memory buffer was used to update the Q-values, and it was re-introduced to the buffer along with its predecessors with its updated priority. This process was performed iteratively until the buffer was empty (Alg. 2). Q-values were computed using *γ* = 0.9.

In line with human decision making combining goal-directed and random exploration [50, 51], action selection took place using an epsilon-greedy approach, with random actions selected with probability *ε* = 0.05, and otherwise the action with the highest *C*(*s, a*)-value (Eq. 2) was selected in every state. The full ERA algorithm is detailed in Alg. 3.

## 3. Results

In this paper we aim to establish the Epistemic Replay Algorithm (ERA) as an efficient reinforcement learning solution, organically resolving the exploration-exploitation trade-off in a biologically satisfying manner; improving on the reward rates of previous algorithms in an environment with multiple rewarded sites while exhibiting replay properties reminiscent of biological agents (Tab. 2). Accordingly, the results are broken down into 6 consecutive sections (Tab. 3), testing for efficiency (Sec. 3.1, 3.2), adaptability (Sec. 3.3) and replay structure (Sec. 3.4, 3.5, 3.6).

**Table 3:** The five experimental setups. The research questions correspond to the 6 hypotheses tested, divided into 3 categories (efficiency [yellow], adaptability [green] and structure [blue]). They are as follows: curiosity can drive efficient exploration; information-seeking is more optimal than uncertainty-seeking; curiosity plays a key role in adapting to change in the reward structure; curiosity promotes the replay of branching points; curiosity contributes to realistic replay patterns as observed in neuroscience experiments (Tab. 2); curiosity is a satisfying explanation for vicarious trial and error (VTE) [15]; and curiosity can enable quick adaptive rerouting in face of an obstacle. The variable *w_i_ ∈* ℝ is the weight of the epistemic component in the computation of the C-values; *r_i_* stands for epistemic reward type, which can be uncertainty-based (abs) or information-based (diff); and *walls* stands for the possibility to bump into walls.

| Sec | Environment | Question | Variable | Range |
| --- | --- | --- | --- | --- |
| 3.1 | Simple linear maze | Exploration | $w_i$ | 0, 2, ..., 10 |
| 3.2 | Flickering task | Information vs uncertainty | $r_i$<br>$w_i$ | *abs, diff*<br>0, 2, 10 |
| 3.3 | Non-stationary maze | Adaptation to reward change | $w_i$ | 0, 2, ..., 10 |
| 3.4 |  | Branching points & curiosity |  |  |
| 3.5 | Restricted double-T maze | Replay patterns | $w_i$ | 0, 2, ..., 10 |
| 3.6 | Non-restricted double-T maze | Branching points & VTE | *walls*<br>$w_i$ | *yes, no*<br>0, 2, 10 |
| 3.7 | Restricted double-T maze | Adaptation to obstacles | — | — |

### 3.1 Curiosity enables efficient exploration

In the first experiment, we showed the relevance of incorporating curiosity and curiosity-driven replay into the study of reinforcement learning. To do so, a simple linear maze was devised (Fig. 2A) consisting of a straight corridor with the start location at the southernmost end point, a large reward (*r* = 5) at the distal (northernmost) end, and a small proximal “decoy” reward (*r* = 0.5) in the southern third of the maze. Each episode consisted of 25 steps, meaning that an agent that is maximally greedy with respect to the proximal reward could achieve a cumulative reward rate of 10 each episode, while one that is greedy with respect to the distal reward would achieve a cumulative reward rate of 50.

**Figure 2:**
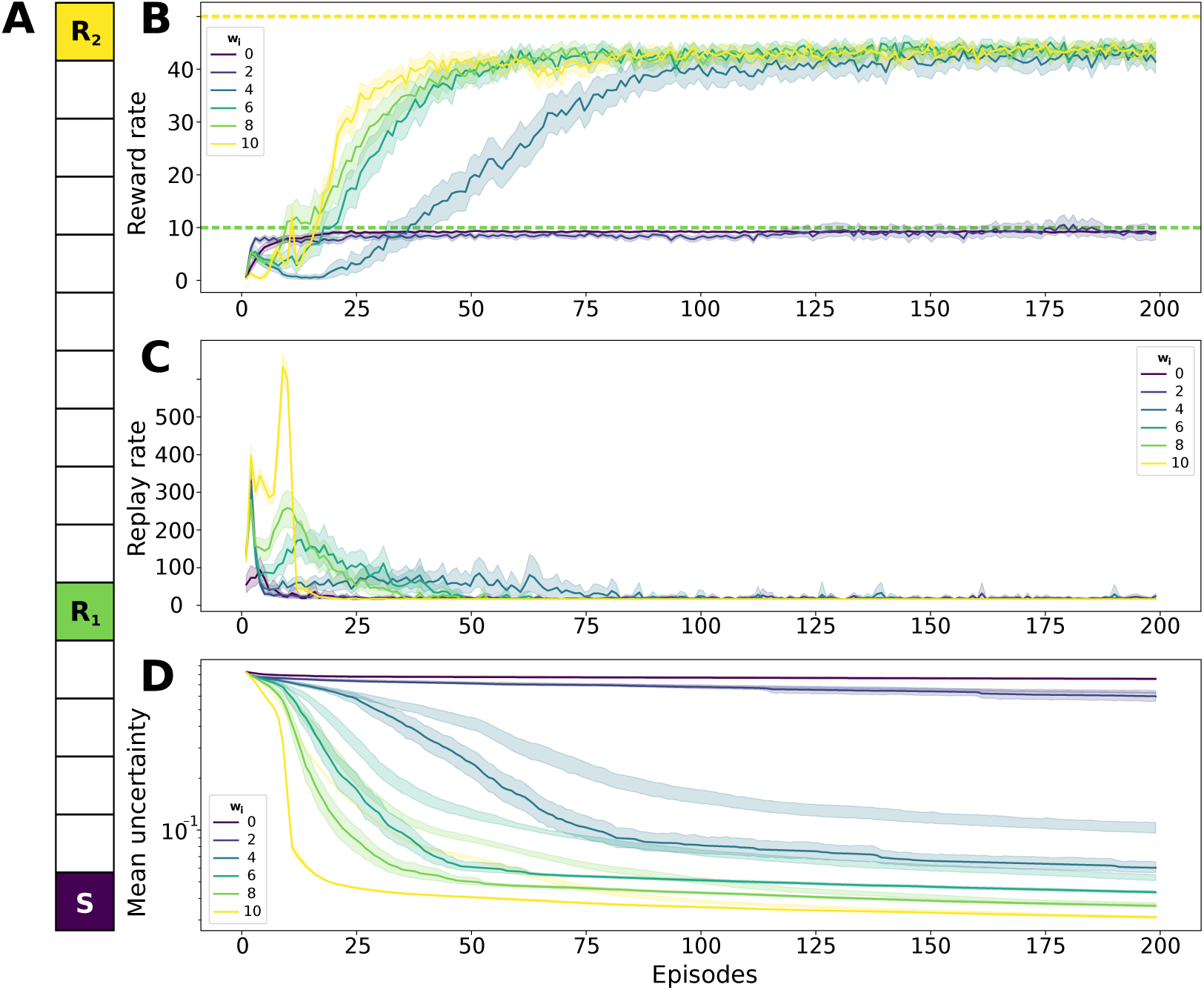
**A** The linear maze. *S*: starting location of each episode; *R*_1_: small proximal reward; *R*_2_: large distal reward. **B** Cumulative reward per episode for 6 different agents. *w_i_*: the weight of the epistemic utility (*Q^i^*-) values (expected information gain about reward uncertainty), where 0 means no curiosity and 10 means that the agent is 10-times more attracted to epistemic rewards than to food rewards. The dashed lines show the theoretical reward rates of the distal (top) and proximal (bottom) rewards. **C** The number of replay steps performed over each episode. **D** The temporal evolution of the mean of the maximal *Q^i^*-value over all states.

As the agent needed to cross the location of the proximal reward to reach the distal one, an overly exploitative algorithm was bound to choose to stay and exploit the decoy (to receive the reward, the agent needed to deliberately choose the “stay” action in the rewarded state). In order to counteract the attractive effect of the proximal reward, exploration needed to be prioritized over exploitation. Six agents were tested in this environment with varying degrees of curiosity (*w_i_ ∈ {*0, 2, 4, 6, 8, 10*}*), where the non-curious (*w_i_* = 0) agent is virtually identical to the hippocampal replay model of Massi et al. [18], and is thus considered the control model.

According to our results, the presence of curiosity modeled by the *Q^i^*-values significantly and consistently contributed to higher reward rates. The reward rate of the non-curious agent converged relatively quickly to that corresponding to the exploitation of the small proximal rewarded site (Fig. 2B). While an only marginally curious agent (*w_i_* = 2) performed at around the same level with slightly quicker convergence; adding further motivation for information seeking (*w_i_ ≥* 4) gradually transformed the behavior. The more curious an agent was, the sooner it achieved a high and stable reward-rate close to that of an ideal agent exploiting the large distal reward. This increasing optimality is due to two factors: first, being more information-motivated meant that the agent explored more quickly, thus allowing it to find the large distal reward sooner; and second, more curious agents replayed epistemically attractive states more often, enabling a more rapidly diminishing epistemic reward, and a more quickly converging extended Q-table that guides behavior. Consequently, during the initial phase of the task (10-30 episodes) curious agents preferred exploring their surroundings (even if this yielded sub-optimal reward rates), but then this drive gradually disappeared and the agent switched its focus to the rewards. This behavior is similar to that obtained by an optimistic initialization of the *Q^f^* -values; however, our solution uses realistic estimates of rewards at all times, and can adapt to stochasticity in a way classical optimistic initialization would not permit (Sec. 3.3).

During the initial phase, replay is more prevalent (Fig. 2C) which is in line with previous findings in rodents [10, 47]. This holds especially true with the more curious agents, as they simply have higher C-values present all across the maze. This spike in replay activity quickly drops with exposure, which is a phenomenon widely observed in biology. [48]

To characterize overall uncertainty, we opted for computing the mean of the maximal *Q^i^* over the entire state space; that is, we calculated the action with the highest *Q^i^* in each state and then averaged all the values computed this way. Plotting this measure over each episode (Fig. 2D) yields dynamics inversely correlated to the reward rates, that is to say that the less curious an agent, the more uncertainty remains in its model, and thus the less reliable its final policy, mostly due to a lack of exploration. In contrast, more curious agents tend to forage farther in the corridor: we have observed a gradual shift in the most distal state explored with increasing *w_i_*. After *w_i_* = 6, the distal reward was fairly consistently reached within the first 100 episodes, however, the more curious an agent was, the sooner it reached the northernmost end of the corridor, proving the adequacy of curiosity in both exploration and replay.

### 3.2 Curiosity is robust to irreducible uncertainty

As illustrated in the previous section, curiosity is a viable tool for enabling informed and efficient exploration; however, the formalization of this drive is not trivial. Using absolute uncertainty (i.e. the total variance of the reward-function) as a drive, while adequately reproducing certain aspects of curious behavior, would fall victim to the attractive effect of inherently uncertain transitions in the state space. Thus, in our model of curiosity, we proposed and tested an agent attracted to the absolute *difference* in model uncertainty. This latter quantity will be high in unknown states, but will gradually diminish with exposure irrespective of the nature of the observed events, deterministic or otherwise. In this sense, epistemic rewards can be considered as a measure of “informativeness” of a given observation, generating a transient drive for information in the early phases of task execution, striking a naturally emerging balance between early exploration and late exploitation.

We modified the Linear Maze task (Sec. 3.1), placing the agent in the corridor in a way that it had to choose between either going to the south to exploit a proximal but stochastic reward (*r* = 3, *q* = 0.5); or to the north where a distal but constant reward (*r* = 4, *q* = 1) was placed (Fig. 3A). Using episodes of 25 steps, the deterministic reward would yield a theoretical reward rate of 60, while the proximal reward rate would be 30. We tested 5 agents in this maze: one non-curious; a pair of total uncertainty-seeking (curiosity directed towards absolute uncertainty); and a pair of information-seeking agents (curiosity directed towards the detected difference in uncertainty, corresponding to our implementation). Both pairs consisted of a mildly-(*w_i_* = 2) and a highly-(*w_i_* = 10) curious member. In order to enable the recognition of stationary stochasticity, the time window over which the model was calculated was increased to *n* = 10.

**Figure 3:**
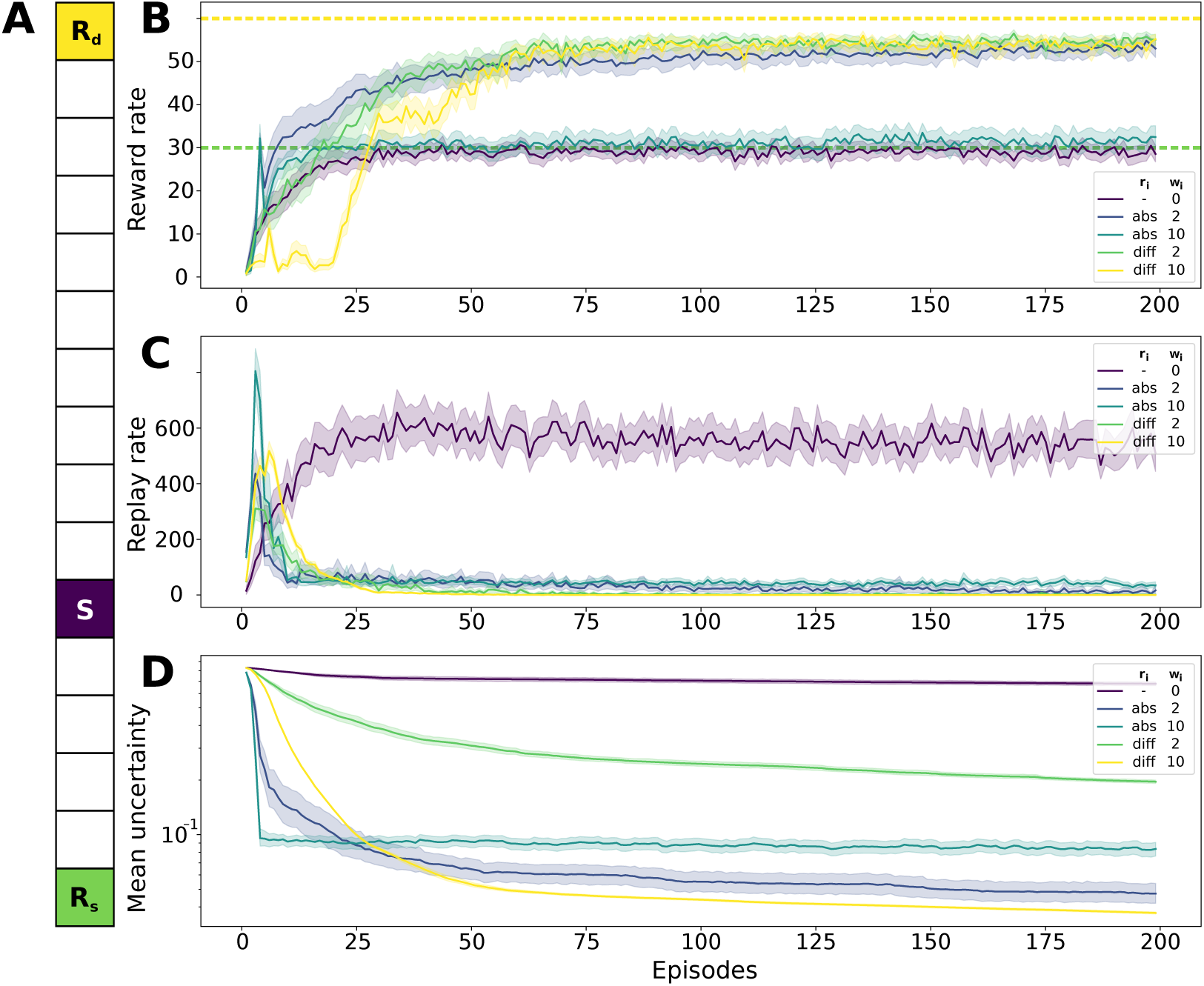
**A** The linear maze with the stochastic reward. *S*: starting location of each episode; *R_s_*: stochastic proximal reward; *R_d_*: deterministic distal reward. **B** Cumulative reward per episode for 6 different agents. *r_i_*: the type of epistemic reward used, where *abs* means absolute uncertainty (”uncertainty-seeking” agent), while *diff* means a difference in uncertainty after learning (”information-seeking” agent); *w_i_*: the weight of the epistemic utility (*Q^i^*-) values. The dashed lines show the theoretical reward rates of the distal (top) and proximal (bottom) rewards. **C** The number of replay steps performed over each episode. **D** The temporal evolution of the mean of the maximal *Q^i^*-value over all states.

As per the previous task, the non-curious agent found and exploited the proximal stochastic reward, simply due to its accessibility (Fig. 3B). Conversely, the slightly curious agents (*w_i_*= 2) both found and exploited the distal constant reward. As expected, this is the result of the curiosity-driven exploration leading to the discovery and quick consolidation of the optimal policy. Interestingly, however, the highly curious agents yielded differing reward rates: while the information-seeking agent reached the distal reward, the uncertainty-seeking one performed similarly to the non-curious agent. The behavior of the latter is due to the fact that albeit this algorithm consistently found the distal reward, its *Q^i^*-values associated with the southernmost state never decreased (due to its constant stochasticity, Fig. 3D), overriding the drive for food rewards. This was not the case for the *w_i_* = 2 uncertainty-seeking agent, since the classical *Q^f^* -values associated to the distal reward could still overcome the *Q^i^*-values in the southern end of the maze, even despite the latter remaining relatively high.

This illustrates that while an optimal policy is possible with an uncertainty-seeking agent, it is heavily dependent on parameter selection (*w_i_*), while the information-seeking agent remains highly robust under any level of curiosity.

Regarding computational cost, since replay is initiated by surprise, the non-curious agent produced continuous reactivations over the course of the entire experiment (Fig. 3C). This means that, even though the agent itself is model-based, without an a priori estimate of the model uncertainty, the algorithm is forced to constantly update its representation upon novel observations. On the other hand, all curious agents – regardless of their type – restricted their replay to the initial (model learning) phase, rendering them significantly more efficient. This shows that our curious (information-driven) agent is: (i) more optimal in scenarios necessitating directed exploration than traditional agents; (ii) more optimal in the face of stochasticity than both traditional and uncertainty-seeking agents; (iii) more computationally efficient than traditional agents using replay.

### 3.3 Curiosity allows for adaptation in a changing environment

After having addressed the question of optimality and efficiency, we turned to specific scenarios. Notably, one of the most extensively studied aspects of RL with hippocampal replay is the ability to adapt to non-stationarity. Several studies have proposed different versions of a T-maze, in which the agent learned the location of a reward, which was then consequently displaced, forcing the algorithm to re-explore its environment and establish a new optimal policy. We propose that in a scenario such as this, the directed exploration due to curiosity will allow us to recover the reward rate more quickly.

Accordingly, a T-maze with a small proximal (*r* = 0.5) and a large distal (*r* = 3) reward was devised. After 200 episodes, both rewards disappeared, and a new large distal reward emerged, equidistant from the previous two, and identical in its value to the large distal reward of the first phase. This ensured that no matter the initial policy, all agents would have an equal footing for finding and exploiting the new rewarded location post-change. Episodes in this maze consisted of 30 steps, yielding a theoretical cumulative reward rate of 12 for an agent greedy with respect to the small proximal reward; and a cumulative reward rate of 54 for an agent greedy with respect to the pre- or post-reward change large distal reward. Similarly to the previous experiment, *n* = 10 was selected for the time window of model integration, to allow for a more defined transition period.

Up until the reward change, we observed similar reward-, replay- and *Q^i^*-dynamics to the Linear maze (Sec. 3.1). After episode 200, the reward rates dropped significantly, only recovering in case of a non-zero *w_i_* (Fig. 4B). This recovery was, as observed in rats [44], accompanied by an immediate replay event starting at the moment of reward extinction comparable to the initial replay in its range (see [45, 20]), but this replay rate was inversely proportional to the height of the initial peak and thus, to *w_i_* itself. Consequently, it can be concluded that agents that had previously produced a massive replay event to reduce their uncertainty (curious agents) were able to more efficiently adapt to a later change due to better and quicker convergence of Q-values and more extensive exploration. It is interesting to point out that while the non-curious agent remains stuck at the previous reward location after the reward change (possibly due to the *Q^f^* -values of these states asymptotically approaching but never reaching zero), agent *w_i_* = 2, while unable to consistently reach the large distal reward in the first 200 episodes, was nevertheless capable of achieving similar reward rates to the more curious agents in the second half of the experiment. This means that curiosity does not only promote initial exploration, but is also a mechanism enabling adaptation after a change in the environment by perturbing sub-optimal local maxima in the traditional *Q^f^* -value landscape. When it comes to the mean uncertainty in the maze, a second drop was observed right after the reward change (Fig. 4C). This indicates a second phase of exploration due to the non-stationarity. This auxiliary exploration, however, is notably absent in the non-curious agent, reaffirming that information-seeking is at he basis of adaptive behavior.

**Figure 4:**
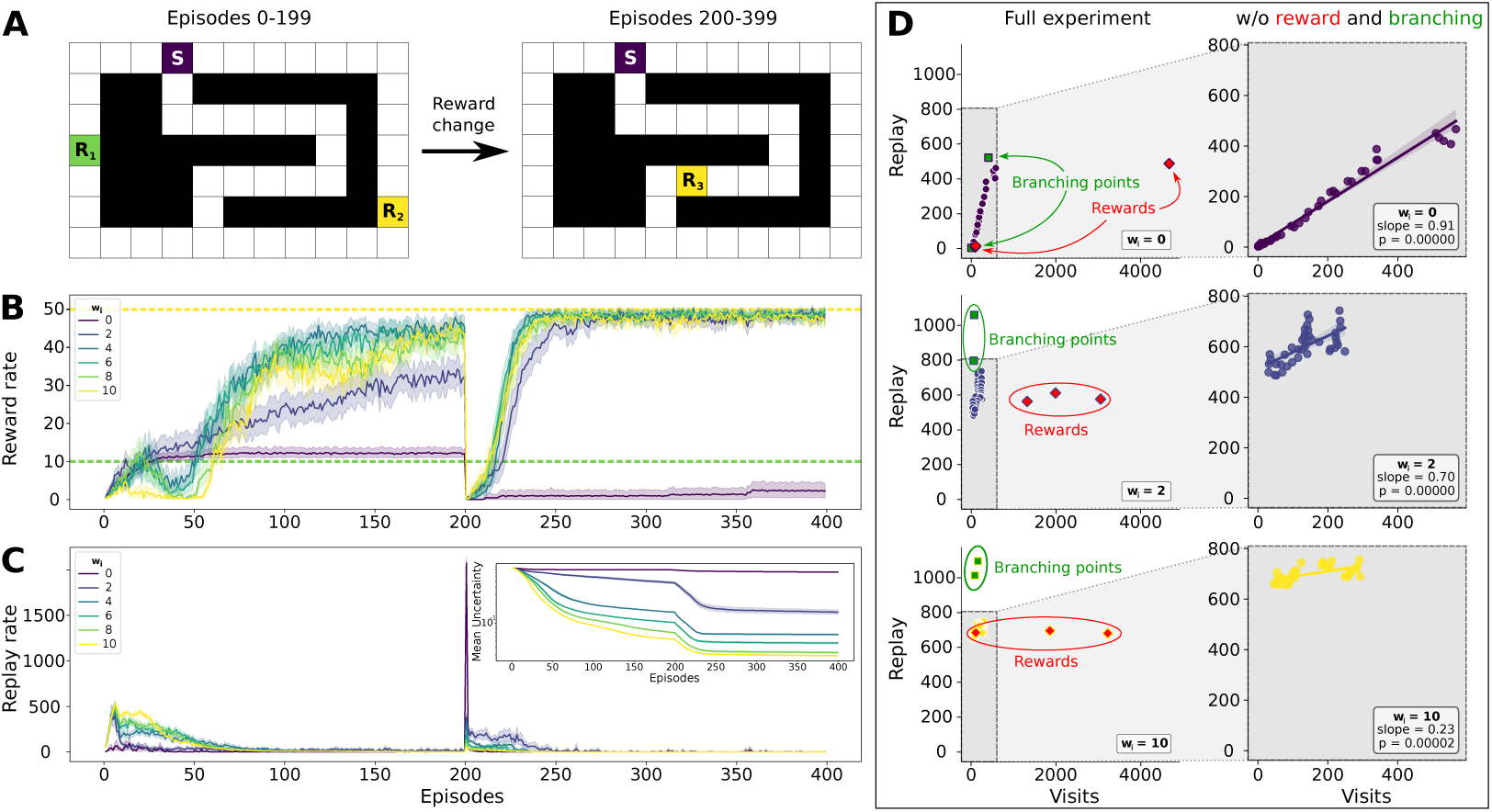
**A** The non-stationary maze. *R*_1_: small proximal reward; *R*_2_: large distal reward. *R*_3_ large distal reward after reward change. **B** Cumulative reward per episode. **C** The number of replay steps performed over each episode. Inset: temporal evolution of the mean of the maximal *Q^i^*-value over all states. **D** The number of replay steps in each state vs the amount of time the agent spent in a given state (visits) for the non-curious, marginally curious (*w_i_* = 2) and extremely curious (*w_i_*= 10) agents. The left column shows the entire state space, while the right column has the reward locations and branching points removed. A linear was fit to each data set in the right column.

### 3.4 Replay is structured around branching points

Having established the Epistemic Replay Algorithm as a robust, adaptive and efficient algorithm with high performance in scenarios necessitating informed exploration, we now turn to its biological relevance. We propose the hypothesis that curiosity and information seeking play a role not only directly in behavior, but also indirectly in learning via awake hippocampal reactivation. In this and the following sections, we will compare the structure and dynamics of our replay towards epistemic rewards with those of classical algorithms or observations from biology. To this end, we will continue to use different implementations of the T-maze, as they are predominantly used in the literature on hippocampal reactivations.

As O’Neill et al. observed in rats [46], replay of a given state is proportional to the amount of time spent in the place field of the corresponding neuron. We observed a similar tendency in the non-stationary maze (Fig. 4D). It appears that there is a linear correlation between the reactivation and the time spent in the given state (Fig. 4D, right), and this trend persists among all levels of curiosity.

This tendency holds true for all except a handful of states (Fig. 4D, left), namely the rewarded sites and branching points. Rewarded sites are, per definition, the states in which an optimal agent spends most of its time; nevertheless, their presence in the replay content is not disproportionally high, as most of the time spent in these states is dedicated to exploiting the rewarded action following the (already converged) policy. Consequently, once the initial surprise is reduced and the model is stabilized, no further inference (replay event) is triggered about these states as no new information is incorporated into the model.

Yet another outlier is branching points. We defined branching points as states from which more actions are available than from an average state of the environment (for further detail see Sec. 3.6). Branching points are visited just as often as other states the agent passes through; however, they are replayed significantly more than the rest of the environment. In the coming sections we aim to shed light onto the mechanism behind this phenomenon.

### 3.5 The spatiotemporal properties of replay resemble awake hippocampal reactivations

To continue the investigation of the replay structure, we needed to adapt the environment such that it resembles previous studies – both in biology and in reinforcement learning – more closely. This meant a serious truncation of the state space and a restriction of movement: awake hippocampal replay has been thoroughly studied in mazes consisting of uni-directional corridors, the double T-maze of Redish et al [44] being of particular interest to us, given its symmetrical composition, 3 branching points of different natures (decision point giving to a dead end, decision point giving to 2 similar corridors, confluence point) and small number of states and actions.

Since our non-curious agent is virtually identical to their solution, and since they did not manage to show elevated stopping in the southernmost branching point, we opted for the maze implemented by Massi et al. [18], which can be readily decomposed into a unidirectional (northbound) central corridor, and two southbound arms. The reward (*r* = 1) is placed in the western arm for the first 100 episodes (where it is delivered upon stepping onto the rewarded state), after which it is moved to the eastern arm. The central corridor connects to the two arms via two branching points (the top one where paths diverge and the bottom one where paths converge); and there exists a third branching point in the middle of the central corridor, giving to a dead end. As the “stay” action was not available in this environment, the agent was forced to continually move forward, and thus episodes were delimited by the agent arriving at the start state.

Given that this maze is solvable without directed curiosity, we did not expect our agent to outperform previously existing solutions. Accordingly, and in line with the literature, the reward rates of all tested agents quickly converged to the optimum (Fig. 5B) where they remained until the reward change. This initial increase was smooth for the non-curious agent, while more noisy for the agents with higher *w_i_* values, indicating the usual initial exploration phase. After the reward change, recovery happened within the first 10 episodes, with the most curious agent lagging slightly behind the others. This could be explained by the fact that this agent performed the least amount of replay after the reward change (Fig. 5C). Just as before, initial replay was more dominant for the more curious agents, and replay after the reward change was more prevalent for the non-curious agent.

**Figure 5:**
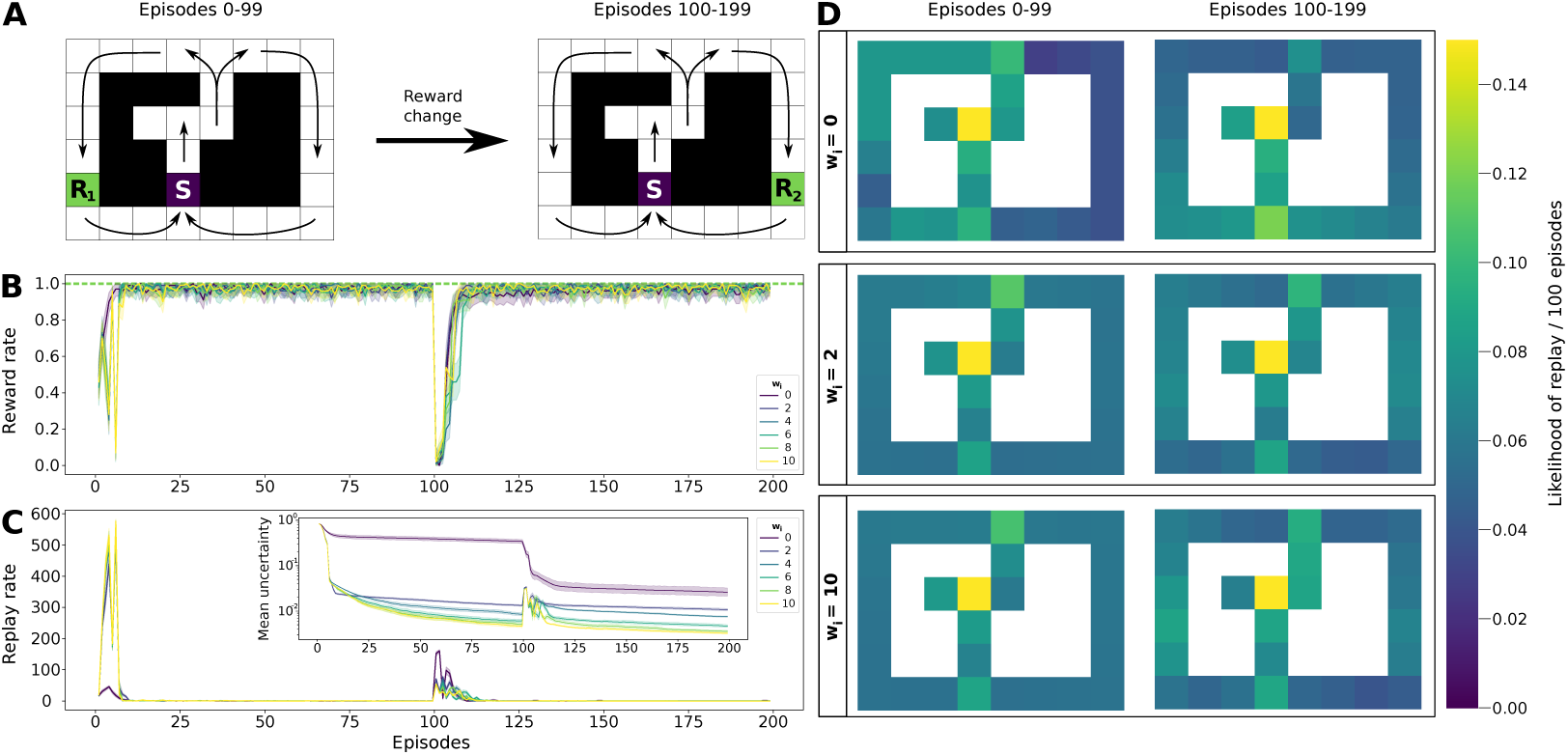
**A** The unidirectional double-T maze. Rewards *R*_1_ and *R*_2_ are of the same value. Arrows show the permitted direction of motion in each corridor. Backtracking is only allowed from the central dead end. **B** Cumulative reward per episode. **C** Number of replay steps performed over each episode. Inset: temporal evolution of the mean of the maximal *Q^i^*-value over all states. **D** The likelihood of each state being replayed, decomposed into 100 episodes pre- (1st column) and post reward change (second column). The top represents the the non-curious-, the middle row the marginally curious-(*w_i_* = 2) and the bottom row the extremely curious (*w_i_* = 10) agents.

The mean uncertainty in the maze (Fig. 5C inset) followed a similar pattern as in the case of the non-stationary maze (Sec. 3.3), however, due to the significantly smaller state space we could observe two novel phenomena. Firstly, even the non-curious agent managed to greatly decrease its own uncertainty due to the baseline exploration ensured by epsilon-greedy decision making. Secondly, within this setup, small-scale changes in uncertainty were still visible upon computing the mean *Q^i^*-values, which explains why we can see a peak of uncertainty after the reward change in the case of the agents. This peak appears as these agents are likely to replay and thus back propagate the discounted reward uncertainty to farther parts of the maze, essentially broadening an initially localized phenomenon. Ultimately, this is the biggest novelty of our method: the ability to back propagate and thus anticipate uncertainty in distant parts of the maze, allowing for the agent to perform decisions informed not only by the rewards, but the delayed potential information gain as well.

As expected, the structure of the replay varied noticeably between the curious and the non-curious agents. While we ascribe most of this difference to the effect of prioritizing the reactivation based on the *||*Δ*C||*_1_-values (containing discounted expected information gain); we would like to address the fact that non-curious agents explore the environment less thoroughly, yielding a potentially smaller memory pool; albeit this effect should be minimal in the double-T maze.

In the initial phase of the experiment (first 100 steps), we observed that while classical replay shows a strong bias towards replaying the rewarded corridor (*p* = 9.216*e −* 05, Mann-Whitney-Wilcoxon rank-sum test [M.W.W from this point onward] on the *w_i_* = 0 agent comparing relative replay of the left and right corridors), this bias is non-significant in case of the most curious agent (M.W.W *p* = 0.47). Indeed, as the initial broad exploration (and replay) phase coincides with the convergence of the classical *Q^f^* -values, no further food-reward focused replay is necessary to establish an optimal policy. This unbiased replay of the two corridors reflects biological evidence observed in a similar environment [13], and could be attributed to a consequence of the reward bias [39, 40, 41, 6, 42, 43], or more precisely a special case of it with respect to the dense epistemic rewards distributed quasi-uniformly over the state space.

Conversely, after the reward change, the curious agent performs a replay event much more focused than that of the classical agent, showing a significant reward bias towards classical food-rewards (linear regression of the relative replay frequencies vs the distance from the reward in case of the *w_i_* = 10 agent: slope = *−*4.4*e −* 3, *p* = 1.49*e −* 162; vs agent *w_i_* = 10: *−*2.9*e −* 3, *p* = 1.28*e −* 144). This change is the result of two different phenomena: firstly, the curious agent explores significantly more in the initial phase of the task, reducing model uncertainty to a minimum even before the reward change, making additional replay superfluous; and secondly, while the disappearing reward only results in a small local negative change in the reward function of the classical agent (taking several trials to back propagate), the curious agent receives an additional epistemic reward at the time of the model update, making it highly motivated to perform focused replay of the missing reward, allowing for quicker forgetting of the old site and quicker learning of the new site. This is in line with the claims of Mattar and Daw [17], who observed that replay has a tendency of progressing from broad and non-specific towards local and specific.

In case of the non-curious agent, this reward bias is missing in the second half of the experiment, replaced by a “starting point bias” (linear regression of the relative replay frequencies vs the distance from the start in case of the *w_i_* = 0 agent: slope = *−*7.1*e −* 3, *p* = 6.35*e −* 232). This latter likely emerges due to the frequent visits of the starting location due to the task framework and explained by O’Neill et al. [46]. It is important to distinguish this phenomenon from the “initiation bias” [9, 10, 38], or the fact that online replay is often centered around the agent’s current state. This latter is a natural consequence of our algorithm, as replay is only triggered once the replay buffer is non-empty due to a significant *local* change in Q-values.

Nevertheless, the reward and start point biases are not the only effect structuring replay. Branching points seem to be overly represented in the replay across all agents and all phases of the experiment, although the non-curious agent shows significantly more bias towards them in the first half (M.W.W. for the first and second halves of the experiment: *p* = 5.005*e −* 3 vs *p* = 1.473*e −* 2 for the non-curious; *p* = 4.449*e −* 4 vs *p* = 8.899*e −* 4 for the most curious).

While an elevated replay *of* the decision points is yet to be tested in foraging rodents, an elevated rate of *stopping* in decision points is indeed well documented [44, 13]. But what if this behavior is not actually triggered by an internal understanding of “decisions”, but by a more primitive mechanism? We propose that an elevated replay around *branching points* might be a result of the internal world model and curiosity: essentially states with a higher-than-average number of predecessors and successors are likely to reactivate more often and are learned at a different rate than the rest of the world given their elevated information content. This effect might have been overlooked in previous literature, as the starting point of foraging often coincided with the confluence of the different paths the animal can take, resulting in the “starting point bias” overshadowing the “branching point bias”.

### 3.6 Preferential stopping in branching points is explained by curiosity

In order to test our hypothesis regarding the nature of the branching points, we needed a setup in which we could manipulate the number of available actions in each state. Accordingly, two versions of the Double-T maze were considered (Fig. 6B): one where all actions were permitted across all states (*permissive* condition), and one where bumping into walls was forbidden (*restrictive* condition), thus promoting 3 states of the maze to branching points. It is important to note here, however, that even the *restrictive* version of this environment was more permissive than the original Double-T maze: in these tests, movement direction and staying in a given state were allowed, thus an episode length of 25 needed to be defined once again before the agent was forcefully returned to the start. Just as before, we used a reward of *r* = 1 (now delivered upon selecting to “stay”) at reward site *R*_1_, which was displaced after 100 steps (reward site *R*_2_; Fig. 6A). In both environments, we tested 6 agents of varying levels of curiosity.

**Figure 6:**
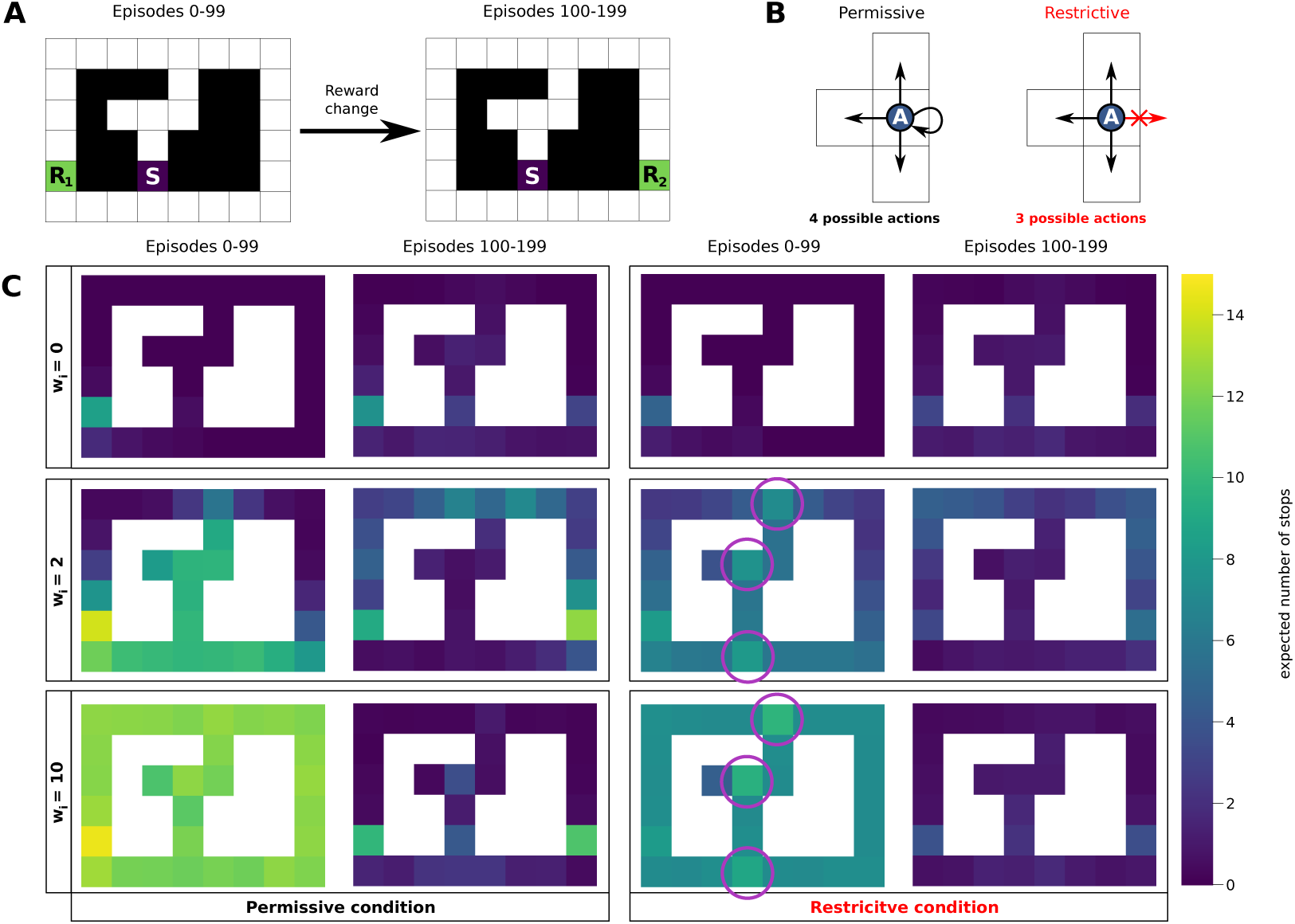
**A** The double-T maze. *R*_1_ and *R*_2_ are of the same value. **B** The *”permissive”* and *”restricitve”* conditions. In the former case the agent always perceives 4 possible actions, while in the latter only those actions are perceived that could result in physical motion. **C** The expected number of stops performed by the agent in each state over the course of the first and second 100 episodes of the task. The two left-hand columns represent the permissive condition; while the two right-hand columns represent the restrictive condition, both decomposed into before- and after reward change respectively. The 3 rows represent 3 different levels of curiosity.

Further, as it was already pointed out in the previous section, there is little to no data on the decision/branching point bias in the replay content; however, there is a very clear tendency of pausing in these points to perform replay (vicarious trial and error [15]). Consequently, in this section we will focus on the number of behavioral pauses the agent performed in different parts of the maze.

Irrespective of the experimental condition, we observed a similar tendency in the behavior as those observed in the replay structure as before: the non-curious agent almost exclusively stops at and around the reward locations both before and after the reward change, as these are the states where large reward prediction errors are encountered. In contrast, the more curious agents pause more uniformly all over the state space. These “exploratory” stops tend to take place closer to the start location in the early experiment, and propagate further away in time as the agent visits more of its environment, a phenomenon most clearly observed in case of the *w_i_* = 2 agent (Fig. 6C).

Crucially, however, a bias towards pausing in “branching points” only emerged under two coinciding conditions: firstly, only curious agents stopped preferentially in branching points (M.W.W. of branching points vs. the rest of the state space in the first 100 trials under the restrictive condition: *p* = 0.784 [*w_i_* = 0] vs *p* = 5.434*e −* 3 [*w_i_* = 10]); secondly, this preferential stopping only took place in case of the *restrictive* condition (M.W.W. of branching points vs. the rest of the state space in the first 100 trials for the *w_i_* = 10 agent: *p* = 0.5244 [permissive] vs *p* = 5.434*e −* 3 [restrictive]).

These findings seem to point in the direction of curiosity playing a role in vicarious trial and error. If the restrictive condition in and of itself had been sufficient to trigger the branching point bias, one could argue that this behavior exclusively emerges due to the model-based nature of replay: more predecessor and successor states means more replay sweeps converging in the given state, resulting in more replay independently of curiosity. And while this is true for the replay *content* (Fig. 5D), we have seen that this phenomenon is not enough to explain behavioral pauses. We propose that an important contributing factor to preferential stopping in branching points is the excess information which can act as an epistemic reward to the agent. Consequently, we promote the idea that no inherent understanding of “decision points” is needed in order to reproduce vicarious trial and error-type behavior in these points.

### 3.7 Curiosity boosts adaptation to new obstacles

At the time of writing this article, one of the best known models of hippocampal replay in reinforcement learning is that of Mattar and Daw [17]. While they primarily treat replay as an optimal way to accelerate learning, their prioritized sweeping-type algorithm has recently been expanded by Antonov and Dayan [31] to incorporate replay in the face of uncertainty. Both of these normative accounts of hippocampal replay involve the computation of a replay priority over the experienced state-action pairs in a prioritized sweeping-type algorithm. This priority can be decomposed into a product of a Gain and a Need term, where Gain is proportional to a predicted 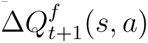 (*s, a*) weighted by the policy, while Need is defined as the expected discounted number of visits to a given state in the environment.

This is the framework that was expanded by Antonov and Dayan into a world consisting of probabilistic belief states in order to account for uncertainty. Their *Exploratory Gain* encodes the expected local improvement of the policy incorporating the potential information gain after learning; while their *Exploratory Need* term reflects the likelihood of encountering any given potential update belief state according to the current prior beliefs. Using this framework, it is indeed possible to generate replay events extending to-wards uncertain state transitions, resulting in policies promoting targeted exploration.

As opposed to their solution, which builds on a normative framework of optimal action selection, we position ourselves as a heuristic account of hippocampal replay in the face of uncertainty, proposing an easily expandable and efficient reinforcement learning algorithm of low computational complexity. While the two algorithms are similarly attracted to novelty in unexplored environments, to offer a qualitative account of the differences between these works, we recreated the Tolman-maze [52] experiment analyzed by Antonov and Dayan, wherein the introduction of a wall forces the agent to reroute its policy towards the rewarded location (Fig. 7). We compared their agent with our maximally curious ERA-algorithm (*w_i_*= 10). As environmental uncertainty was now a result of a change in the transition model, we opted to compute the *Q^i^*-values as a discounted expected absolute change in the uncertainty (entropy) of the *transition*-function, instead of the reward function. Further, to render learning more visually apparent, we opted for a decreased replay threshold (*θ* = 0.005 *×* (*w_f_* + *w_i_*)).

**Figure 7:**
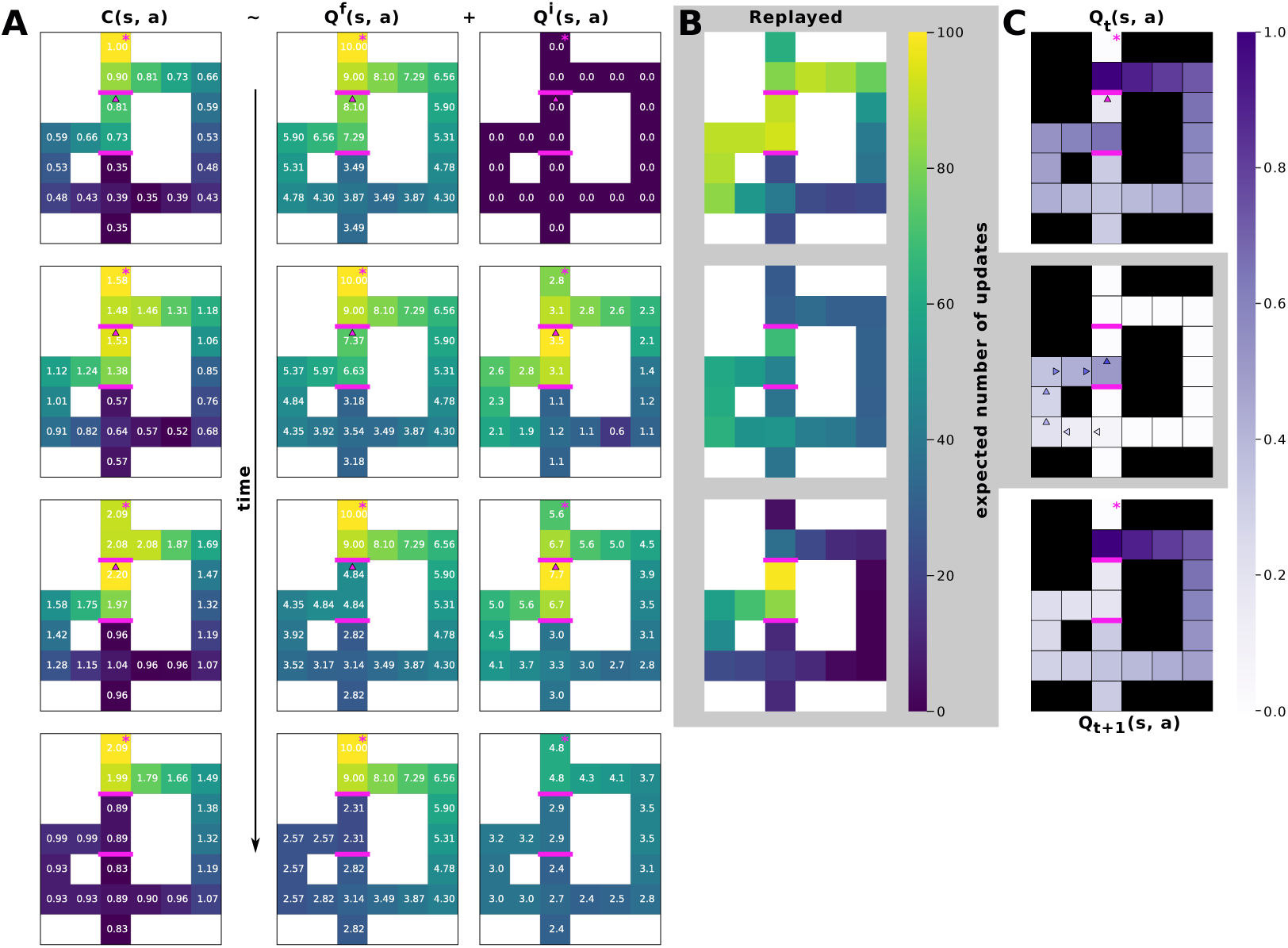
**A** The temporal evolution of the *C*(*s, a*) (first column) and the corresponding *Q^f^* (*s, a*) and *Q^i^*(*s, a*)-values (second and third column respectively) over the course of 3 replay steps after the discovery of a wall. Walls are denoted in magenta, the rewarded state is marked with an asterisk. The first row depicts the initial state, in which the agent was forced to bump into a newly introduced wall (this action is indicated by a magenta triangle). The second row shows the same values after one replay step. Here the agent was forced to perform the same action (magenta triangle) as before. This process was repeated until the values converged. **B** The average number of replay steps over the maze after each action. **C** The same experiment performed by Antonov and Dayan [31]. The first row represents the Q-values before the discovery of the wall, the second row is the elicited replay event (arrows indicate the replayed actions and the color represents temporality), and the third row depicts the Q-values after the replay. The figure is directly reproduced using the code published alongside their article.

One important liberty we took regarding the experiment was the removal of the Northern wall in the initial phase of learning: our agent was pretrained on an environment without any blockages. This is important, as the model of Antonov and Dayan is capable of producing a replay event towards potentially uncertain parts of the maze (the location where walls may or may not be) even if change has never been observed – assuming that the *possibility* of change is encoded in the belief state. This results in occasional policy changes promoting the active (re-)discovery of potentially uncertain areas of the maze. In practical terms, this can be observed for example as a spontaneous replay event towards a previously encountered wall. While the vanilla version of ERA only triggers replay upon directly encountering uncertainty, we could achieve a similar result by implementing a forgetting mechanism: a continuous additive bonus on the *Q^i^*-values proportional to the amount of time passed since the last visit to a given state-action pair. Nonetheless, we consider this work to be beyond the scope of this paper.

Once the initial policy established, walls are introduced to the maze, and the agent is forced to bump into the one closer to the rewarded state from the South. In case of both models, this change will elicit a small decrease in the *Q^f^* -values, but this change is usually too small for a proper replay event to take place. In case of the ERA-algorithm, however, this experience is accompanied by a local *increase* of the *Q^i^*-values, resulting in a sizable reactivation, propagating this change to further states (Fig. 7A-B). Upon further experience confirming the presence of the wall, these replay events will elicit a full local reevaluation of the Q-values, achieving total convergence after 3 replay iterations, or once the existence of the wall is estimated to have a probability of *p* = 1.

Given the same setup, the vanilla model of Antonov and Dayan produces no significant reactivation, forcing trial-and error unlearning of the local Q-values. In order to achieve quick convergence, *Sequence Replay* was introduced, a mechanism promoting the replay of chains of actions of pre-defined length leading up to the state-action couple of interest. This extension of their algorithm achieves a similar result to ERA (Fig. 7C). The authors remain transparent about the arbitrary nature of this solution.

## 4. Discussion

We realized a reinforcement learning algorithm, ERA, equipped with prioritized sweeping, and a two-dimensional Q-table that captures both extrinsic rewards (e.g., food) and expected discounted information gain, thereby providing a minimal operationalization of curiosity. Replay was produced as a function of these Q-vectors, and the direct and indirect effects of these reactivations were systematically evaluated across five different experimental setups, designed as epistemic foraging tasks [53], where uncertainty-driven information gathering directly shapes policy optimization.

It was shown that curiosity enables agents to efficiently explore and find hidden rewards, even in the presence of decoys, while exploitation of said rewards emerges naturally as the uncertainty of the internal model decreases. Importantly, this effect was significantly more robust in the case of agents showing a secondary drive towards information gain, but not in the case of agents driven by pure uncertainty reduction.

This phenomenon has been addressed in detail by [54, 55, 56]. Essentially, uncertainty can be decomposed into so-called aleatoric (irreducible) and epistemic (reducible) components. Aleatoric uncertainty is a property of the environment connected to its stochasticity, while epistemic uncertainty characterizes the information that can be extracted via observation. A truly random process has high aleatoric uncertainty as the outcome is always surprising, but little epistemic uncertainty, as these outcomes are hardly informative of any underlying structure.

Using the absolute uncertainty (i.e. the total variance of the reward-function) as an epistemic reward is similar to being attracted to aleatoric uncertainty: highly stochastic observations may repeatedly drive exploration even though no informative structure can be learned from them, as illustrated by the “noisy TV problem” described in Burda et al. [57]. Positing uncertainty, thus, as a reward function would correspond less to a model of curiosity and more to a model of surprise-seeking, irrespective of its potential to inform decision making. In contrast, using the absolute difference in model uncertainty as epistemic reward ensures that, given a stationary (stochastic or otherwise) process, the epistemic reward observed by the agent always converges to zero after *n* visits, resulting in diminishing returns. Therefore, only states with non-stationarity could remain epistemically attractive to the agent, in line with our definition of curiosity, thus overcoming the noisy TV problem.

Accordingly, ERA agents showed significant adaptability in the case of non-stationary environments: we observed early, more exhaustive exploration and replay, yielding quicker, more focused learning after a change in the environment structure. This result is in line with findings in the study of spatial navigation in rodents, reproducing several key aspects of the spatiotemporal organization of realistic hippocampal reactivations. Further, we man-aged to reproduce preferential stopping in certain loci that we call branching points, showing that an underlying non-homogeneous representation of the environment and active information seeking are necessary and sufficient for the emergence of said activity. In other words, no further understanding of the task outcomes is required for the agent to preferentially stop to replay in these “decision points”, a concept which often appears in the literature [44, 13].

One key aspect of ERA is the natural emergence of a balance between exploration and exploitation. Indeed, initial phases of both behavior and replay are driven by a strong bias towards informative states; this bias, however, naturally diminishes with exposure, giving rise to the exploitation of the most optimal policy. However, unlike in the case of optimistic initialization, ERA will show an increase in curiosity following changes in the environment, triggering directed (re-)exploration. This way, ERA comes as a naturally emerging solution to the exploration-exploitation dilemma.

The central component of our algorithm is that of replay. Hippocampal replay has been shown to contribute to learning, memory consolidation, and planning. Evidence suggests that its structure might be more than a direct reactivation of experience, with a wide range of studies narrowing down its spatiotemporal organization. One leading hypothesis from the field of Reinforcement Learning proposes that replayed sequences are likely driven by a back-propagated priority value related to rewards [17, 19]. Here we propose that this priority value might incorporate further dimensions, most notably that of information gain, in line with Active Inference Theory [24].

A novel algorithm unifying hippocampal replay, epistemic rewards and reinforcement learning is that of Antonov and Dayan [31]. And while both Antonov and Dayan’s method and the ERA algorithm behave similarly in the face of novel environments (ie., promoting the directed exploration of unknown states and actions), their predictions differ significantly in the case of stochastic changes in a well-known state space. While the model of Antonov and Dayan can propose pre-play towards estimated uncertainty through a forgetting-like mechanism, the ERA-algorithm’s strength is using uncertainty as a “carrying signal” to back propagate *Q^f^* -values in case of an unexpected change. We believe that biological agents clearly exhibit both of these phenomena (vicarious trial and error and one-shot unlearning), and the two methods show how curiosity might contribute to efficient learning in the face of different forms of uncertainty.

It is worth noting that reactivation patterns generated by the two algorithms differ slightly, reproducing different phenomena observed in the study of hippocampal replay. While the vanilla algorithm of Antonov and Dayan can perform sequential (backwards) replay back-propagating from perceived uncertainty, replay steps remain unitary after learning, and only using the *Sequence Replay* heuristic can they be linked up into backwards sequences. While the ERA algorithm currently produces no pre-play, its replay patterns show several properties in line with biology. Following the definitions of Gupta et al. [6], replay produced by ERA is mostly “unordered”, spreading concentrically from rewarding (epistemically or otherwise) states, nevertheless with clear “backward”, “forward” and “imaginary” patterns appearing in restricted areas of the maze. Building on these computational results, we would like to propose that hippocampal replay – similarly to ERA – may actually follow a breadth-first search like reactivation pattern, one which, observed in narrow, linear environments, can give rise to the more well-studied depth-first like replays.

Finally, while replay in case of the algorithm of Antonov and Dayan is launched at pre-defined points in time, the ERA algorithm uses an adaptive replay, suspending behavior when re-evaluation of its model is necessary. Importantly, the structure of the behavior produced this way is in line with research on rodents [15, 44, 13].

In conclusion, we hypothesize that curiosity plays an important role in both spatial navigation and hippocampal replay in the context of epistemic foraging. Curiosity, as we modeled it, could be considered as the drive towards epistemic rewards, just as hunger is the drive towards food rewards. One advantage of this formalism is its infinite expandability – namely that any number of additional needs could be added to the algorithm, essentially creating a pyramid of needs [58], where each one is satisfied by a different set of policies among which the agent can freely switch in accordance with its current necessities and possibilities.

Indeed, assuming *k* needs satisfied by rewards *{r*_1_*, …r_k_}*, and preferences *{w*_1_*, …w_k_}*, we could formalize the *subjective* reward value at any given point as a scalar product between the reward vector and the preference vector, that is 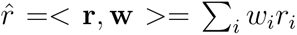. In this case, the subjective Q-value at any moment could be computed as:

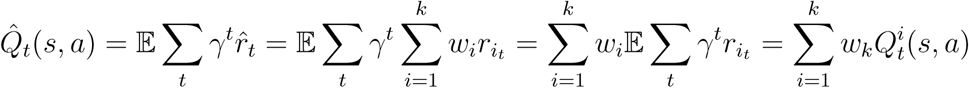

where 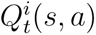 is the *need-specific* Q-value of state *s* and action *a* given need *i*, meaning that is it is computed as if the policy was only optimized for this singular need. Given this formalism, it is entirely possible to compute each *need-specific* dimension of the Q-table via classical RL-algorithms, and use their linear combination (the *subjective* Q-value) for the establishment of the current policy. One such system can utilize the full (objective) reward for value learning, irrespective of its momentary preferences; and any change of this latter can provoke an immediate change in policy without the necessity for unlearning or re-learning. Additionally, the computational complexity of one such system is linear in the number of needs modeled.

Consequently, balancing hunger, curiosity, and potential further homeostatic needs is an essential next step for bridging intrinsically motivated reinforcement learning with homeostatic control [59, 60, 61]. In the scope of this work and for the sake of simplicity, we opted for using a linear combination of the *Q^f^* and *Q^i^*-values, transforming rewards into a common currency [62, 63, 37]. This, in essence, produces results similar to methods trained on a linear combination of food- and epistemic rewards right away, given the static weights of the agent’s preferences. However, since our framework keeps track of the separate reward dimensions, it is theoretically possible not only to combine the Q-values, but also to train a meta-agent to tune the weights of the linear combination, or even arbitrate between the available dimensions, yielding a high-level allostatic RL system [64, 65, 66].

## Acknowledgments

The authors would like to thank Oscar Guerrero-Rosado, Denis Sheynikhovich and Paul F.M.J. Verschure for useful discussions. This work was supported by the European Commission’s CAVAA Project (EIC 101071178), the ANR ELSA project (ANR-21-CE33-0019) and the Neuro-Flex project (ANR-24-CE37-5256-02).

## References

[1] H. F. Ólafsdóttir, D. Bush, C. Barry, The role of hippocampal replay in memory and planning, Current Biology 28 (2018) R37–R50. URL: https://www.cell.com/current-biology/fulltext/S0960-9822(17)31441-0. 10.1016/j.cub.2017.10.073.

[2] A. Forli, W. Fan, K. K. Qi, M. M. Yartsev, Replay and representation dynamics in the hippocampus of freely flying bats, Nature (2025) 1–3. URL: https://www.nature.com/articles/s41586-025-09341-z. 10.1038/s41586-025-09341-z.

[3] A. C. Schapiro, E. A. McDevitt, T. T. Rogers, S. C. Mednick, K. A. Norman, Human hippocampal replay during rest prioritizes weakly learned information and predicts memory performance, Nature Communications 9 (2018). 10.1038/s41467-018-06213-1.

[4] T. K. Leonard, K. L. Hoffman, Sharp-wave ripples in primates are enhanced near remembered visual objects, Current biology: CB 27 (2017) 257–262. URL: https://www.ncbi.nlm.nih.gov/pubmed/28041797. 10.1016/j.cub.2016.11.027.

[5] J. ÓKeefe, J. Dostrovsky, The hippocampus as a spatial map. preliminary evidence from unit activity in the freely-moving rat, Brain Research 34 (1971) 171–175.

[6] A. S. Gupta, M. A. van der Meer, D. S. Touretzky, A. D. Redish, Hippocampal replay is not a simple function of experience, Neuron 65 (2010) 695–705. 10.1016/j.neuron.2010.01.034.

[7] H. F. Ólafsdóttir, D. Bush, C. Barry, The role of hippocampal replay in memory and planning, Current Biology 28 (2018) R37–R50.

[8] M. A. Wilson, B. L. McNaughton, Reactivation of hippocampal ensemble memories during sleep, Science 265 (1994) 676–679.

[9] K. Diba, G. Buzsáki, Forward and reverse hippocampal place-cell sequences during ripples, Nature Neuroscience 10 (2007) 1241–1242. 10.1038/nn1961.

[10] D. J. Foster, M. A. Wilson, Reverse replay of behavioural sequences in hippocampal place cells during the awake state, Nature 440 (2006) 680–683. 10.1038/nature04587.

[11] G. Girardeau, K. Benchenane, S. I. Wiener, G. Buzsáki, M. B. Zugaro, Selective suppression of hippocampal ripples impairs spatial memory, Nature neuroscience 12 (2009) 1222–1223.

[12] V. Ego-Stengel, M. A. Wilson, Disruption of ripple-associated hip-pocampal activity during rest impairs spatial learning in the rat, Hip-pocampus 20 (2010) 1–10.

[13] A. Johnson, A. D. Redish, Neural ensembles in ca3 transiently encode paths forward of the animal at a decision point, Journal of Neuroscience 27 (2007) 12176–12189. 10.1523/jneurosci.3761-07.2007.

[14] A. D. Redish, Mental time travel: A retrospective, Hippocampus 35 (2024). 10.1002/hipo.23661.

[15] E. C. Tolman, Prediction of vicarious trial and error by means of the schematic sowbug., Psychological Review 46 (1939) 318–336. 10.1037/h0057054.

[16] H.-T. Chen, M. A. A. van der Meer, Paradoxical replay can protect contextual task representations from destructive interference when experience is unbalanced, bioRxiv (Cold Spring Harbor Laboratory) (2024). 10.1101/2024.05.09.593332.

[17] M. G. Mattar, N. D. Daw, Prioritized memory access explains planning and hippocampal replay, Nature Neuroscience 21 (2018) 1609–1617. 10.1038/s41593-018-0232-z.

[18] E. Massi, J. Barthélemy, J. Mailly, R. Dromnelle, J. Canitrot, E. Poniatowski, B. Girard, M. Khamassi, Model-based and model-free replay mechanisms for reinforcement learning in neurorobotics, Frontiers in Neurorobotics 16 (2022). 10.3389/fnbot.2022.864380.

[19] R. Caźe, M. Khamassi, L. Aubin, B. Girard, Hippocampal replays under the scrutiny of reinforcement learning models, Journal of neurophysiology 120 (2018) 2877–2896.

[20] M. Khamassi, B. Girard, Modeling awake hippocampal reactivations with model-based bidirectional search, Biological cybernetics 114 (2020) 231–248. 10.1007/s00422-020-00817-x.

[21] K. L. Stachenfeld, M. M. Botvinick, S. J. Gershman, The hippocampus as a predictive map, Nature neuroscience 20 (2017) 1643–1653.

[22] G. Pezzulo, E. Cartoni, F. Rigoli, L. Pio-Lopez, K. Friston, Active inference, epistemic value, and vicarious trial and error, Learning & Memory 23 (2016) 322–338.

[23] R. Kaplan, K. J. Friston, Planning and navigation as active inference, Biological cybernetics 112 (2018) 323–343.

[24] T. Van de Maele, B. Dhoedt, T. Verbelen, G. Pezzulo, A hierarchical active inference model of spatial alternation tasks and the hippocampal-prefrontal circuit, Nature Communications 15 (2024) 9892.

[25] J. Peng, R. J. Williams, Efficient learning and planning within the dyna framework, Adaptive Behavior 1 (1993) 437–454. doi:10.1177/105971239300100403.

[26] A. W. Moore, C. G. Atkeson, Prioritized sweeping: Reinforcement learning with less data and less time, Machine Learning 13 (1993) 103–130. 10.1007/bf00993104.

[27] A. A. Carey, Y. Tanaka, M. A. A. van der Meer, Reward revaluation biases hippocampal replay content away from the preferred outcome, Nature Neuroscience 22 (2019) 1450–1459. 10.1038/s41593-019-0464-6.

[28] A. K. Gillespie, D. A. Astudillo, E. L. Denovellis, D. F. Liu, D. B. Kastner, M. Coulter, D. Roumis, U. T. Eden, L. M. Frank, Hippocampal replay reflects specific past experiences rather than a plan for subsequent choice, Neuron 109 (2021) 3149–3163.e6. doi:10.1016/j.neuron.2021.07.029.

[29] R. S. Sutton, A. Barto, Reinforcement learning: An introduction, The Mit Press, Cambridge, Ma ; London, 2018.

[30] H. Van Seijen, R. S. Sutton, Efficient planning in mdps by small backups, in: Proceedings of the 30th International Conference on International Conference on Machine Learning - Volume 28, ICML’13, JMLR.org, 2013, p. III–361–III–369.

[31] G. Antonov, P. Dayan, Exploring replay, Nature Communications 16 (2025). 10.1038/s41467-025-56731-y.

[32] L. Aubin, M. Khamassi, B. Girard, Prioritized sweeping neural dynaq with multiple predecessors, and hippocampal replays, Lecture notes in computer science (2018) 16–27. 10.1007/978-3-319-95972-6_4.

[33] M. J. Gruber, B. D. Gelman, C. Ranganath, States of curiosity modulate hippocampus-dependent learning via the dopaminergic circuit, Neuron 84 (2014) 486–496. URL: http://orca.cf.ac.uk/96033/1/uncorrected_proof_Gruber_Curiosity_main_text.pdf. doi:10.1016/j.neuron.2014.08.060.

[34] M. J. Gruber, C. Ranganath, How curiosity enhances hippocampus-dependent memory: The prediction, appraisal, curiosity, and exploration (pace) framework, Trends in Cognitive Sciences 23 (2019) 1014–1025. 10.1016/j.tics.2019.10.003.

[35] P. Schwartenbeck, J. Passecker, T. U. Hauser, T. H. FitzGerald, M. Kronbichler, K. J. Friston, Computational mechanisms of curiosity and goal-directed exploration, eLife 8 (2019) e41703. URL: https://elifesciences.org/articles/41703. doi:10.7554/eLife.41703.

[36] E. S. Bromberg-Martin, I. E. Monosov, Neural circuitry of information seeking, Current Opinion in Behavioral Sciences 35 (2020) 62–70.

[37] D. J. Levy, P. W. Glimcher, The root of all value: a neural common currency for choice, Current Opinion in Neurobiology 22 (2012) 1027–1038. 10.1016/j.conb.2012.06.001.

[38] T. J. Davidson, F. Kloosterman, M. A. Wilson, Hippocampal replay of extended experience, Neuron 63 (2009) 497–507. doi:10.1016/j.neuron.2009.07.027.

[39] B. E. Pfeiffer, D. J. Foster, Hippocampal place-cell sequences depict future paths to remembered goals, Nature 497 (2013) 74–79. doi:10.1038/nature12112.

[40] D. Dupret, J. O’Neill, B. Pleydell-Bouverie, J. Csicsvari, The reorganization and reactivation of hippocampal maps predict spatial memory performance, Nature Neuroscience 13 (2010) 995–1002. doi:10.1038/nn.2599.

[41] H. F. Ólafsdóttir, C. Barry, A. B. Saleem, D. Hassabis, H. J. Spiers, Hippocampal place cells construct reward related sequences through unexplored space, eLife 4 (2015). 10.7554/elife.06063.

[42] A. E. Papale, M. C. Zielinski, L. M. Frank, S. P. Jadhav, A. D. Redish, Interplay between hippocampal sharp-wave-ripple events and vicarious trial and error behaviors in decision making, Neuron 92 (2016) 975–982. 10.1016/j.neuron.2016.10.028.

[43] F. Michon, J.-J. Sun, C. Y. Kim, D. Ciliberti, F. Kloosterman, Post-learning hippocampal replay selectively reinforces spatial memory for highly rewarded locations, Current Biology 29 (2019) 1436–1444.e5. 10.1016/j.cub.2019.03.048.

[44] A. D. Redish, Vicarious trial and error, Nature Reviews Neuroscience 17 (2016) 147–159. 10.1038/nrn.2015.30.

[45] S. Palminteri, G. Lefebvre, E. J. Kilford, S.-J. Blake-more, Confirmation bias in human reinforcement learning: Evidence from counterfactual feedback processing., PLoS Computational Biology 13 (2017) e1005684. URL: https://doaj.org/article/7442ee695a624c0d82015d8d9c21227f. 10.1371/journal.pcbi.1005684.

[46] J. O’Neill, T. J. Senior, K. Allen, J. R. Huxter, J. Csicsvari, Reactivation of experience-dependent cell assembly patterns in the hippocampus, Nature Neuroscience 11 (2008) 209–215. 10.1038/nn2037.

[47] S. Cheng, L. M. Frank, New experiences enhance coordinated neural activity in the hippocampus, Neuron 57 (2008) 303–313. doi:10.1016/j.neuron.2007.11.035.

[48] L. Buhyr, A. H. Azizi, S. Cheng, S. Université, Reactivation, replay, and preplay: How it might all fit together, Neural Plasticity 2011 (2011). 10.1155/2011/203462.

[49] F. Trovo, S. Paladino, M. Restelli, N. Gatti, Sliding-window thompson sampling for non-stationary settings, Journal of Artificial Intelligence Research 68 (2020) 311–364. doi:10.1613/jair.1.11407.

[50] S. J. Gershman, Deconstructing the human algorithms for exploration, Cognition 173 (2018) 34–42. 10.1016/j.cognition.2017.12.014.

[51] R. C. Wilson, A. Geana, J. M. White, E. A. Ludvig, J. D. Cohen, Humans use directed and random exploration to solve the explore–exploit dilemma., Journal of Experimental Psychology: General 143 (2014) 2074–2081. 10.1037/a0038199.

[52] E. C. Tolman, Cognitive maps in rats and men., Psychological review 55 (1948) 189.

[53] T. Parr, K. J. Friston, Uncertainty, epistemics and active inference, Journal of The Royal Society Interface 14 (2017) 20170376. doi:10.1098/rsif.2017.0376.

[54] S. C. Hora, Aleatory and epistemic uncertainty in probability elicitation with an example from hazardous waste management, Reliability Engineering & System Safety 54 (1996) 217–223. doi:10.1016/s0951-8320(96)00077-4.

[55] G. Velentzas, C. S. Tzafestas, M. Khamassi, Memory development with heteroskedastic bayesian last layer probabilistic deep neural networks, Workshop on World Models and Predictive Coding in Cognitive Robotics at 2023 IEEE/RSJ International Conference on Intelligent Robots and Systems (IROS 2023) (2023). URL: https://hal.sorbonne-universite.fr/hal-04249890v1. doi:https://hal.sorbonne-universite.fr/hal-04249890.

[56] J. Y. Angela, P. Dayan, Uncertainty, neuromodulation, and attention, Neuron 46 (2005) 681–692.

[57] Y. Burda, H. Edwards, A. Storkey, O. Klimov, Exploration by random network distillation, arXiv:1810.12894 [cs, stat] (2018). URL: https://arxiv.org/abs/1810.12894.

[58] A. Maslow, Theory of Human Motivation, volume 50, Wilder Publications, 1943.

[59] N. Chentanez, A. Barto, S. Singh, Intrinsically motivated reinforcement learning, Advances in neural information processing systems 17 (2004).

[60] P.-Y. Oudeyer, F. Kaplan, What is intrinsic motivation? a typology of computational approaches, Frontiers in neurorobotics 1 (2007) 108.

[61] M. Keramati, B. Gutkin, Homeostatic reinforcement learning for integrating reward collection and physiological stability, eLife 3 (2014). 10.7554/elife.04811.

[62] L. P. Sugrue, G. S. Corrado, W. T. Newsome, Choosing the greater of two goods: neural currencies for valuation and decision making, Nature Reviews Neuroscience 6 (2005) 363–375. URL: https://www.nature.com/articles/nrn1666. 10.1038/nrn1666.

[63] E. S. Bromberg-Martin, M. Matsumoto, O. Hikosaka, Dopamine in motivational control: rewarding, aversive, and alerting, Neuron 68 (2010) 815–834.

[64] P. Sterling, Allostasis: A model of predictive regulation, Physiology & Behavior 106 (2012) 5–15. 10.1016/j.physbeh.2011.06.004.

[65] J. Schulkin, P. Sterling, Allostasis: A brain-centered, predictive mode of physiological regulation, Trends in Neurosciences 42 (2019) 740–752. 10.1016/j.tins.2019.07.010.

[66] O. G. Rosado, A. F. Amil, I. T. Freire, P. F. M. J. Verschure, Drive competition underlies effective allostatic orchestration, Frontiers in Robotics and AI 9 (2022). 10.3389/frobt.2022.1052998.

